# Released Bacterial ATP Shapes Local and Systemic Inflammation during Abdominal Sepsis

**DOI:** 10.1101/2024.03.07.583973

**Authors:** Daniel Spari, Annina Schmid, Daniel Sánchez-Taltavull, Shaira Murugan, Keely Keller, Nadia Ennaciri, Lilian Salm, Deborah Stroka, Guido Beldi

**Author notes:** Corresponding author: Guido Beldi; Freiburgstrasse 18, 3010 Bern.

## Abstract

Sepsis causes millions of deaths per year worldwide and is a current global health priority declared by the WHO. Sepsis-related deaths are a result of dysregulated inflammatory immune responses indicating the need to develop strategies to target inflammation. An important mediator of inflammation is extracellular adenosine triphosphate (ATP) that is secreted by inflamed host cells and tissues, and also by bacteria in a strain-specific and growth phase-dependent manner. Here, we investigated the mechanisms by which bacteria release ATP. Using genetic mutant strains of *Escherichia coli* (*E. coli*), we demonstrate that ATP release is dependent on ATP synthase within the inner bacterial membrane. In addition, impaired integrity of the outer bacterial membrane and bacterial death notably contribute to ATP release. In a mouse model of abdominal sepsis, local effects of bacterial ATP were analysed using a transformed *E. coli* bearing an arabinose-inducible periplasmic apyrase hydrolyzing ATP to be released. Abrogating bacterial ATP release shows that bacterial ATP suppresses local immune responses, resulting in reduced neutrophil counts and impaired survival. In addition, bacterial ATP has systemic effects via its transport in outer membrane vesicles (OMV). ATP-loaded OMV are quickly distributed throughout the body and upregulated expression of genes activating degranulation in neutrophils, potentially contributing to the exacerbation of sepsis severity. This study reveals mechanisms of bacterial ATP release and its local and systemic roles in sepsis pathogenesis.

## Introduction

Worldwide, 11 million sepsis-related deaths were reported in 2017, which accounted for an estimated 19.7% of all global deaths (Rudd et al., 2020). Given its high incidence and immense socio-economic burden, the World Health Organization (WHO) has declared sepsis as a global health priority (Reinhart et al., 2017).

Sepsis is defined as life-threatening organ dysfunction caused by an imbalanced host immune response to infection. Antibiotic treatment remains the main approach to treat sepsis; however, despite the use of broad-spectrum antibiotics, lethality remains high. Immune-modulating strategies are an additional approach to antibiotics to curb excessive inflammation and to support an effective defense against the infectious agents. Until recently however, most trials inhibiting cytokine responses or toll-like receptors failed, indicating that alternative approaches for immunomodulation are required (Cao et al., 2023).

It has been shown that adenosine triphosphate (ATP), as soon as it is secreted into the extracellular space, critically modulates inflammatory and immune responses (Di Virgilio et al., 2020; Eltzschig et al., 2012) by activating ionotropic P2X and metabotropic P2Y receptors (Burnstock, 2020; Junger, 2011). Also, such purinergic signaling critically alters immune responses during sepsis (Dosch et al., 2019; Ledderose et al., 2016; Spari & Beldi, 2020). In particular, we have described that the connexin-dependent secretion of ATP by macrophages initiates an autocrine loop of over-activation, resulting in altered local and systemic cytokine secretion, which exacerbates abdominal sepsis (Dosch et al., 2019). Such sepsis-promoting effects of host-derived extracellular ATP are secondary to inflammation initiated by bacterial pathogens.

Recently, it has been discovered that also bacteria release ATP into the extracellular space (Mempin et al., 2013). Such ATP release might be a conserved mechanism of protection from host defense and precede host responses. ATP release has been shown for a variety of bacteria including the sepsis-associated *Escherichia coli* (*E. coli*) and *Klebsiella pneumoniae* (*K. pneumoniae*) from the Proteobacteria phylum or *Enterococcus faecalis* (*E. faecalis*) and *Staphylococcus aureus* (*S. aureus)* from the Firmicutes phylum (Diekema et al., 2019; Hironaka et al., 2013; Iwase et al., 2010; Mempin et al., 2013; MureJan et al., 2018).

The mechanisms by which inflammatory and immune responses in the host are modulated by such released bacterial ATP have just begun to be elucidated (Spari & Beldi, 2020). In colonized compartments such as the intestine, it has been shown that ATP released by mutualistic bacteria modulates local cellular and secretory immune responses (Atarashi et al., 2008; Perruzza et al., 2017; Proietti et al., 2019a) and in the mouth, bacterial ATP release leads to biofilm dispersal and periodontitis (Ding et al., 2016; Ding & Tan, 2016). However, the role of ATP released by bacteria invading non-colonized compartments, such as the abdominal cavity or the blood in the context of local and systemic infections, remains to be determined.

In this study, we investigated if ATP released from bacteria influences the outcome of abdominal sepsis. We first isolated sepsis-associated bacteria and measured the amount of ATP they release. Second, we analysed the importance of ATP generation at the inner bacterial membrane on ATP release over time in *E. coli*, using *E. coli* with mutations in integral respiratory chain proteins (Mempin et al., 2013). Third, the function of the outer bacterial membrane on ATP release during exponential growth was assessed using porin mutants (Alvarez et al., 2017; Choi & Lee, 2019). Fourth, we investigated local effects of ATP in the abdominal cavity. Lastly, based on the finding that bacteria secrete outer membrane vesicles (OMV) (Schwechheimer & Kuehn, 2015), which contain ATP (Alvarez et al., 2017; Pérez-Cruz et al., 2015), we investigated systemic consequences of OMV-derived bacterial ATP.

## Results

### *E. coli*, one of the major pathogens in sepsis, releases ATP in a growth phase-dependent and strain-specific manner

To assess ATP release of sepsis-associated bacteria, abdominal fluid of patients with abdominal sepsis was sampled and an-/aerobically incubated on LB agar plates (Figure 1A). Twenty-five different colonies were randomly picked and analysed by whole 16S-rRNA sanger sequencing, which resulted in twelve different bacterial species (Figure 1B). From these, the four most clinically important sepsis-associated bacteria *E. coli, K. pneumoniae* (both gram^neg^), *E. faecalis* and *S. aureus* (both gram^pos^) (Diekema et al., 2019; MureJan et al., 2018) were further cultivated for experimental studies. We quantified released ATP over time (Figure 1-figure supplement 1) using a luciferin-luciferase-based assay. A growth phase-dependent release of ATP was observed in all species, peaking during exponential growth (Figure 1C). Cumulative amount of released ATP was quantified using the area under the curve (AUC) of released ATP over time (AUC ATP). ATP release was detected across all assessed species (Figure 1D). To model these findings in mice, abdominal sepsis was induced using a standardized cecal ligation and puncture model (Dosch et al., 2019) (Figure1E). Similar to human samples, Proteobacteria and Firmicutes phyla were predominating (Figure 1F, see Figure 1B). In the mouse model, *E. coli* released notably more cumulative ATP compared to *E. faecalis* and *S. aureus* (Figure 1G-H). The bacterial species assessed in humans and mice (*E. coli*, *E. faecalis*, *S. aureus*) differed on the strain level (sequences deposited, see data availability statement) confirming that ATP release is strain-specific (Mempin et al., 2013). In summary, sepsis-associated bacteria release ATP in a growth phase-dependent and strain-specific manner. *E. coli,* which is one of the most frequent facultative pathogens in patients with abdominal sepsis, released the highest amount of ATP isolated from a standardized mouse model of abdominal sepsis.

**Figure 1.**
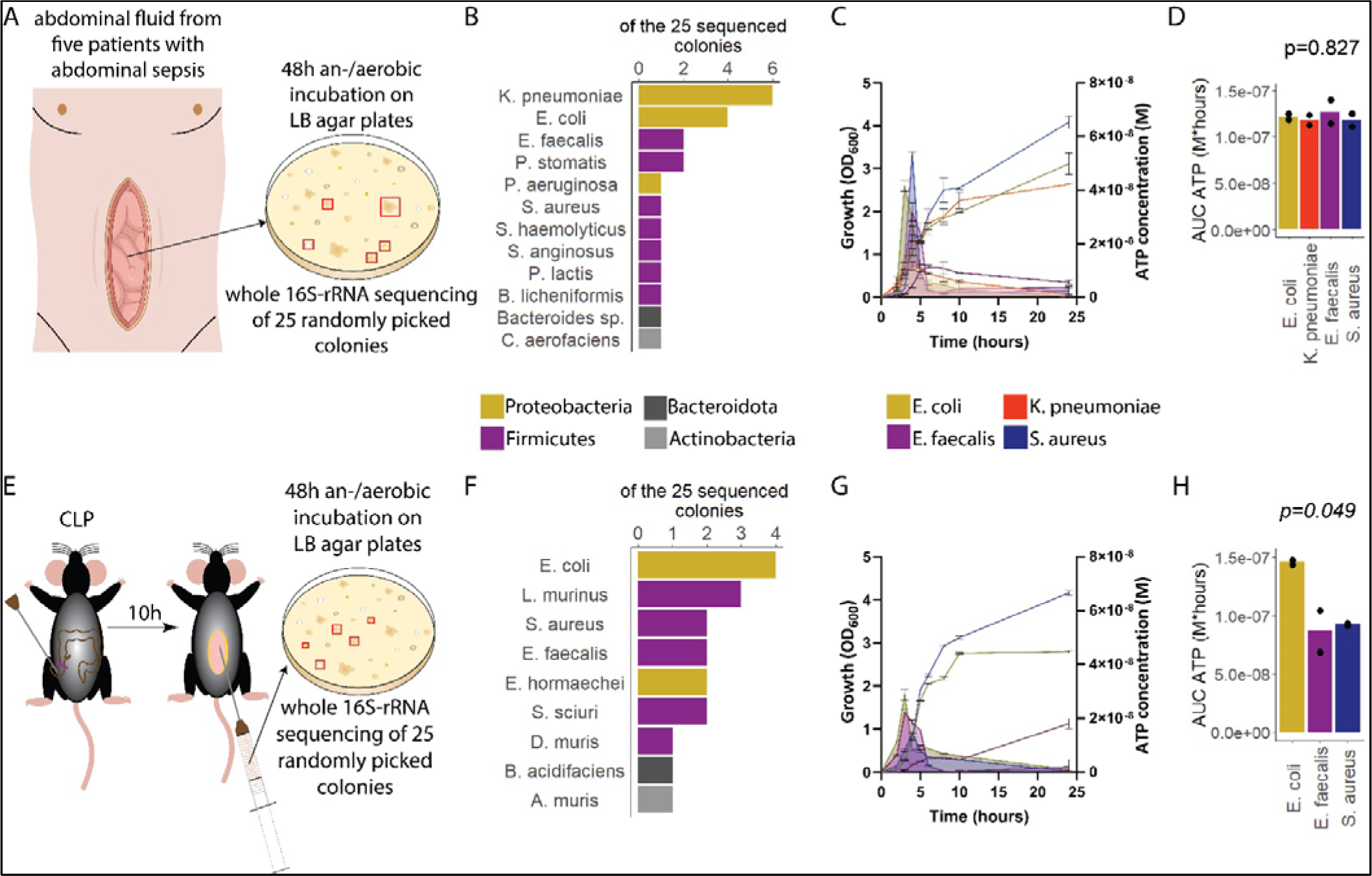
Sepsis-associated bacteria release ATP in a growth phase-dependent manner. **(A)** Experimental approach to isolate and cultivate sepsis-associated bacteria from abdominal fluid of patients with abdominal sepsis. **(B)** Bacterial species identified by whole 16S-rRNA sanger sequencing from abdominal fluid of patients with abdominal sepsis. Three colonies out of 25 could not be identified. **(C)** Measurement of released ATP (M) and growth (OD_600_) over time (hours) from the four sepsis-associated bacteria *E. coli*, *K. pneumoniae*, *E. faecalis* and *S. aureus* isolated from patients. N = 2 independent bacteria cultures. Means and standard deviations are shown. **(D)** Area under the curve (AUC) of released ATP over time (M*hours) of the previously assessed bacteria (cumulative ATP). One-way ANOVA, N = 2 independent bacteria cultures. Means and individual values are shown. **(E)** Experimental approach to isolate and cultivate sepsis-associated bacteria from abdominal fluid of mice with abdominal sepsis. **(F)** Bacterial species identified by whole 16S-rRNA sanger sequencing from abdominal fluid of mice with abdominal sepsis. Seven colonies out of 25 could not be identified. **(G)** Measurement of released ATP (M) and growth (OD_600_) over time (hours) from the three sepsis-associated bacteria *E. coli*, *E. faecalis* and *S. aureus* isolated from mice. N = 2 independent bacteria cultures. Means and standard deviations are shown. **(H)** AUC of released ATP over time (M*hours) of the previously assessed bacteria (cumulative ATP). One-way ANOVA, N = 2 independent bacteria cultures. Means and individual values are shown.

### Bacterial ATP release is dependent on ATP synthesis at the inner bacterial membrane and correlates with bacterial growth

After having demonstrated that sepsis-associated bacteria release ATP, we next questioned whether and how ATP release is dependent on ATP generation. ATP synthase and cytochrome oxidases, which are located in the inner bacterial membrane, are key components of ATP generation under aerobic conditions. Therefore, ATP release of the *E. coli* parental strain (PS) was compared to all available mutants of ATP synthase subunits (Δ*atpA*, Δ*atpB*, Δ*atpC*, Δ*atpD*, Δ*atpE*, Δ*atpF*, Δ*atpH*) and cytochrome *bo3* oxidase subunits (Δ*cyoA*, Δ*cyoB*, Δ*cyoC*, Δ*cyoD*) from the Keio collection (Baba et al., 2006; Yamamoto et al., 2009). The use of different mutants allows to identify the most relevant subunits influencing ATP generation and release. Also it allows to identify a possible correlation of bacterial ATP release with growth, which is a function of ATP generation (Figure 2A) (Galber et al., 2021; Ihssen et al., 2021; Ivančić et al., 2008; Lundin, 2000).

**Figure 2.**
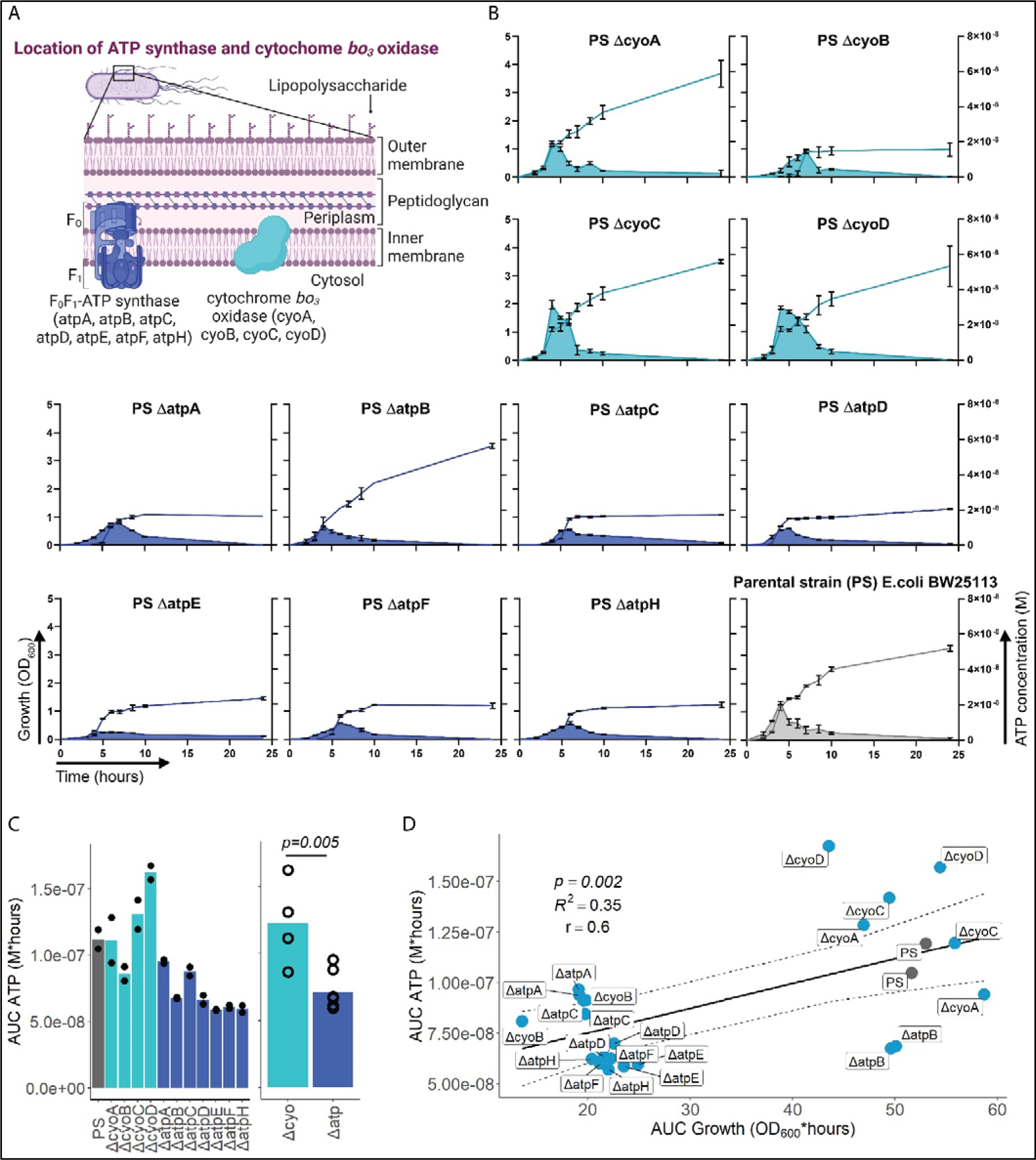
ATP release is dependent on ATP synthesis. **(A)** Illustration depicting the location of ATP synthase and cytochrome *bo_3_* oxidase in gram^neg^ bacteria. **(B)** Measurement of released ATP (M) and growth (OD_600_) over time (hours) from cytochrome *bo_3_*oxidase (cyo) and ATP synthase (atp) mutants. The PS was added as a control. N = 2 independent bacteria cultures. Means and standard deviations are shown. **(C)** Area under the curve (AUC) of released ATP over time (M*hours) of the previously assessed bacteria (cumulative ATP) is shown individually in the left panel. N = 2 independent bacteria cultures. Means and individual values are shown. Means of grouped cyo and atp mutants are compared in the right panel. T-test. Means and individual values are shown. **(D)** Cumulative ATP (M*hours) and cumulative growth (OD_600_*hours) of all assessed cyo and atp mutants and the PS were plotted against each other. Pearson’s correlation (r) and coefficient of determination (*R*^2^) of the applied linear model are depicted. 95% confidence level is shown by the black dashed lines.

Bacterial growth (OD_600_) and ATP release were measured over time and cumulative ATP release (AUC ATP) was assessed (Figure 2B). We first noticed that mutations in subunits of ATP synthase, which is the key enzyme of ATP generation, were generally associated with significantly lower cumulative ATP release compared to mutations in cytochrome *bo3* oxidase subunits (Figure 2C). Therefore, we suspected an interrelation between ATP generation and released ATP (see growth and ATP release curves in Figure 2B). We determined cumulative growth (AUC growth) in addition to cumulative ATP release and indeed, cumulative ATP release and cumulative growth were positively correlated (Figure 2D). Since OD_600_ measurements are most accurate during exponential growth, we additionally assessed released ATP and growth at the individual ATP peak, which was even better positively correlated, strengthening our hypothesis (Figure 2-figure supplement 1). In summary, ATP release is directly dependent on ATP generation at the inner bacterial membrane. Mutations in subunits of bacterial ATP synthase have a higher impact on ATP generation, growth, and ATP release than mutations in subunits of cytochrome *bo3* oxidase (Figure 2C).

### Outer bacterial membrane integrity and bacterial death determine bacterial ATP**LJ**release during exponential growth

We next focused on the outer membrane by challenging its integrity while leaving ATP generation and the inner membrane intact. For that purpose, we used the *E. coli* porin mutants Δ*ompC*, Δ*ompF*, Δ*lamB* and Δ*phoE* (Figure 3A) (Baba et al., 2006; Yamamoto et al., 2009), which have been shown to suffer from impaired outer membrane integrity in varying degrees (Choi & Lee, 2019). ATP release and growth were measured over time including *E. coli* PS as baseline and membrane destabilizing EDTA and stabilizing Ca^2+^ as additional controls (Leive, 1968). Cumulative ATP release (AUC ATP) from the porin mutants were notably different compared to the PS, being lowest in Δ*ompC* and highest in Δ*ompF* (Figure 3B-C).

**Figure 3.**
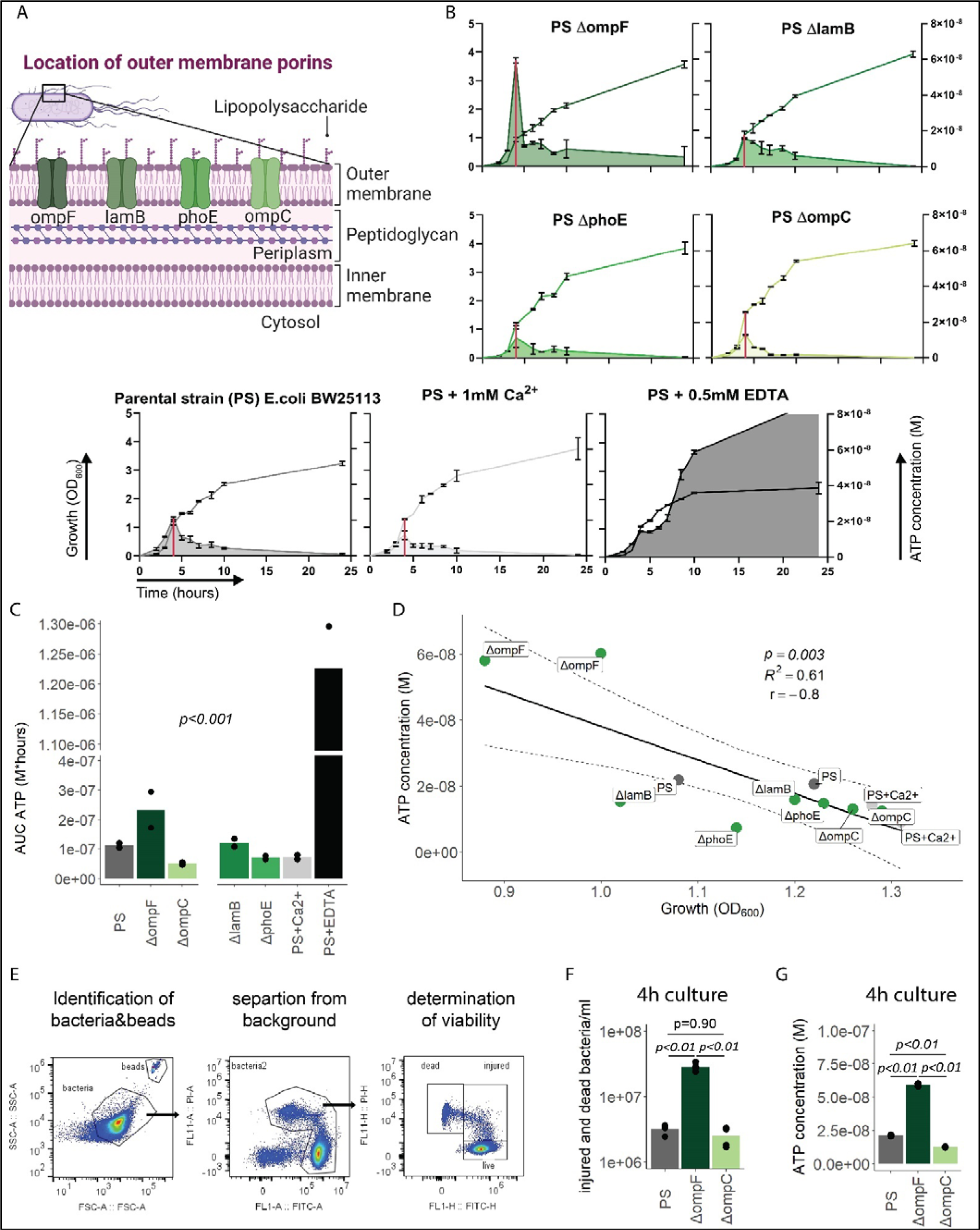
Outer membrane integrity and bacterial death determine bacterial ATPLJrelease during growth. **(A)** Illustration depicting the location of outer membrane porins in gram^neg^ bacteria. **(B)** Measurement of released ATP (M) and growth (OD_600_) over time (hours) from outer membrane porin mutants. The PS and the PS supplemented with either 1mM Ca^2+^ or 0.5mM EDTA were added as controls. N = 2 independent bacteria cultures. Means and standard deviations are shown. The red line marks the individual peak of ATP release and growth (OD_600_) at that time point. **(C)** Area under the curve (AUC) of released ATP over time (M*hours) of the previously assessed bacteria (cumulative ATP). One-way ANOVA, N = 2 independent bacteria cultures. Means and individual values are shown. **(D)** ATP concentration (M) and growth (OD_600_) at the individual peak of ATP release of all assessed outer membrane porin mutants, the PS and the PS+Ca^2+^ (no peak for the EDTA control) were plotted against each other. Pearson’s correlation (r) and coefficient of determination (*R*^2^) of the applied linear model are depicted. 95% confidence level is shown by the black dashed lines. **(E)** Gating strategy to identify added counting beads, live, injured and dead bacteria. **(F)** Quantitative assessment of injured and dead bacteria, as identified by flow cytometry after 4h of culturing (ATP peak) of the PS, Δ*ompF* and Δ*ompC*. One-way ANOVA followed by Tukey post-hoc test, N = 4 independent bacteria cultures. Means and individual values are shown. **(G)** ATP concentration (M) after 4h of culturing (ATP peak) of the PS, Δ*ompF* and Δ*ompC*. One-way ANOVA followed by Tukey post-hoc test, N = 2 independent bacteria cultures. Means and individual values are shown.

Interestingly, the Δ*ompF* mutant had also a very high peak of released ATP (Figure 3B). We hypothesized that this is because of impaired membrane integrity, resulting in ATP release and potentially bacterial death. Thus, we focused on the individual ATP peaks during exponential growth, which were observed after 4h of culturing. There was a strong negative correlation between the individual peak of released ATP (marked by the red line in Figure 3B) and growth at the same time point (Figure 3D).

We did not interfere with ATP generation at the inner membrane but deliberately challenged the outer membrane and tested therewith if destabilization of the outer membrane integrity is associated with bacterial death. Indeed, outer membrane integrity and bacterial death are significantly increased in Δ*ompF* compared to Δ*ompC* and the PS after 4h (ATP peak) of culturing (Figure 3E, F), akin to the amount of released ATP (Figure 3G). We conclude from these data that destabilization of the outer bacterial membrane (as observed with the Δ*ompF* mutant), results in bacterial death that is associated with ATP release.

In summary, outer membrane integrity and finally bacterial death notably contribute to the amount of bacterial ATP release during exponential growth.

### Released bacterial ATP reduces neutrophil counts and impairs survival during abdominal sepsis in mice

Next, we wanted to investigate the function of bacterial ATP release *in vivo*. To study this, we transformed the *E. coli* PS with an arabinose-inducible apyrase (PS+pAPY) and compared it to the PS transformed with the empty vector (PS+pEMPTY) (Proietti et al., 2019b). In this model, ATP released by bacteria is hydrolyzed and consequently depleted by a periplasmic apyrase (Santapaola et al., 2006; Scribano et al., 2014).

Indeed, apyrase induction resulted in a significant reduction of ATP release in PS+pAPY, compared to PS+pEMPTY (Figure 4-figure supplement 1A-B). To test the consequences *in vivo*, apyrase was induced by arabinose three hours before intraabdominal (i.a.) injection into wild type C57Bl/6 mice (Figure 4A). Thereby ATP release was abrogated in the bacteria cultures that were used for injection (Figure 4B). *In vivo*, no difference in ATP levels were detected when ATP was measured directly in the abdominal fluid after four (Figure 4C) and eight hours (Figure 4-figure supplement 1C). This is not surprising given that ATP is rapidly hydrolysed by ectonucleotidases *in vivo* (Eltzschig et al., 2012). After both four (Figure 4D-E) and eight hours (Figure 4-figure supplement 1D-E), no differences in local or systemic bacterial counts were observed. Yet, despite similar bacterial counts, the survival was significantly higher in the absence of bacterial ATP (PS+pAPY) compared to ATP-generating controls (PS+pEMPTY) after i.a. injection (Figure 4F).

**Figure 4.**
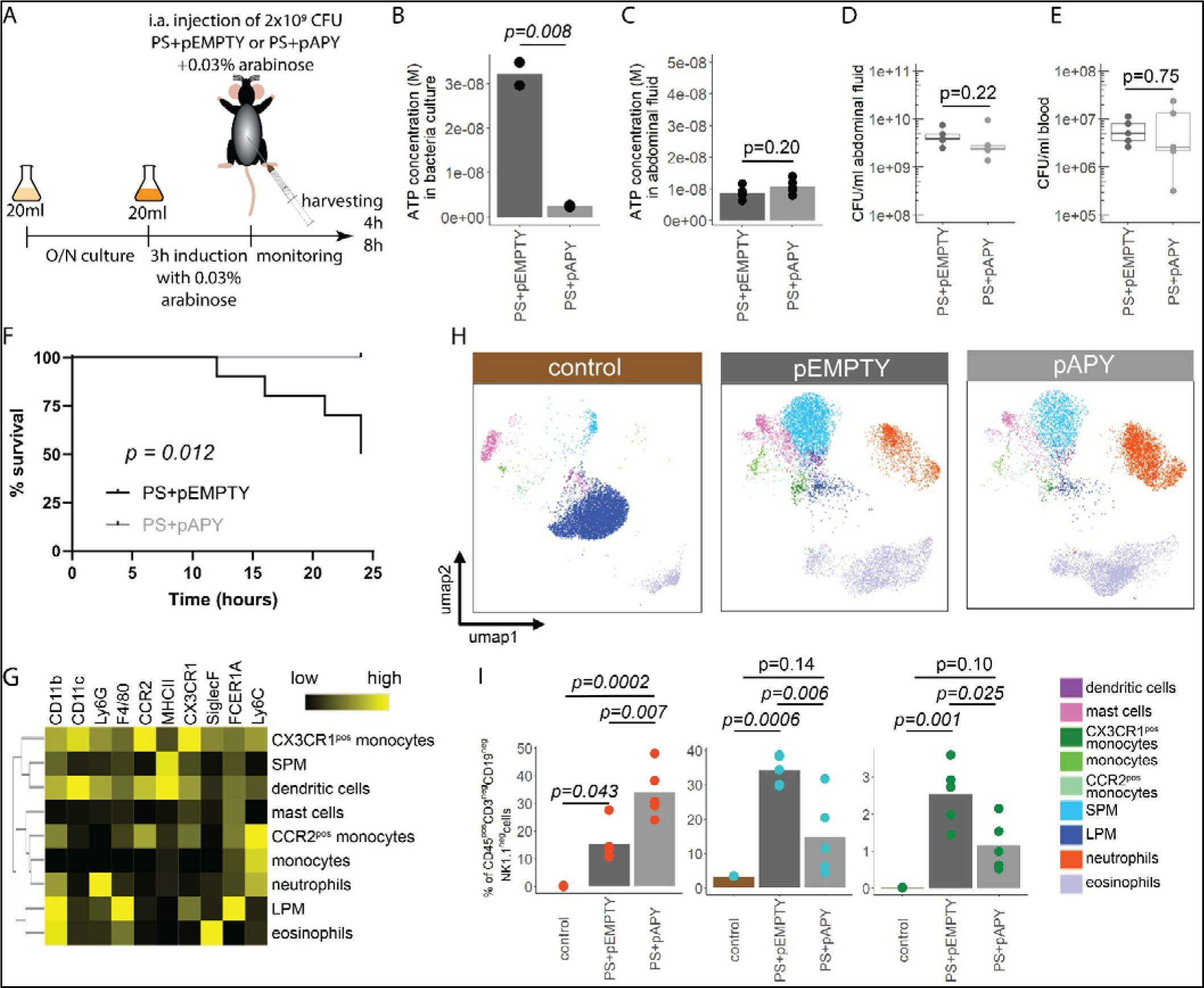
Bacterial ATP reduces neutrophil counts and reduces survival in a mouse model of abdominal sepsis. **(A)** Experimental approach to determine the local role of bacterial ATP *in vivo*, i.a. injecting PS+pEMPTY or PS+pAPY. **(B)** Measurement of released ATP (M) in bacteria culture supernatant immediately before bacteria were i.a. injected. T-test, N = 2 independent bacteria cultures. Means and individual values are shown. **(C)** Measurement of ATP (M) in abdominal fluid from mice four hours after i.a. injection of bacteria. T-test, n = 5 animals per group of N = 2 independent experiments. Means and individual values are shown. **(D)** Quantitative assessment of colony forming units in abdominal fluid and **(E)** blood from mice four hours after i.a. injection of bacteria. Wilcoxon rank sum test, n = 5 animals per group of N = 2 independent experiments. Means and individual values are shown. No growth for controls was detected. **(F)** Kaplan-Meier curves of mice after i.a. injection of bacteria. Log-rank test, n = 10 animals per group. **(G)** Heatmap showing surface marker expression (x-Axis), which was used to characterize the different immune cell populations (y-Axis). **(H)** Concatenated (n = 5 animals for each treatment group, n = 3 animals for control group of N = 2 independent experiments) and down-sampled images of immune cell populations characterized in the abdominal cavity four hours after sham treatment or i.a. injection of bacteria. **(I)** Abundance of neutrophils, small peritoneal macrophages (SPM) and CX3CR1^pos^ monocytes in abdominal fluid from mice four hours after sham treatment or i.a. injection of bacteria. One-way ANOVA followed by Tukey post-hoc test, n = 5 animals for each treatment group, n = 3 animals for control group of N = 2 independent experiments. Means and individual values are shown.

Next, we asked how this difference in bacterial ATP release affects the immune system. Therefore, inflammatory cells in the abdominal cavity were characterized using flow cytometry (Figure 4G). As expected, a disappearance reaction of large peritoneal macrophages (LPM) was observed after both, *E. coli* PS+pEMPTY and *E. coli* PS+pAPY i.a. injection compared to sham controls after four (Figure 4H) and eight hours (Figure 4-figure supplement 1F) (Ghosn et al., 2010). Such LPM disappearance following abdominal *E. coli* infection is a result of free-floating clots composed of LPMs and neutrophils and important for effective pathogen clearance (Salm et al., 2023; Vega-Pérez et al., 2021; Zindel et al., 2021) but not dependent on ATP release according to our data. Interestingly, however, the number of small peritoneal macrophages (SPM) and CX3CR1^pos^ monocytes was significantly reduced, whereas neutrophils were significantly increased almost up to 8 hours (Figure 4I, Figure 4-figure supplement 1G) in bacterial ATP-depleted abdominal sepsis (PS+pAPY). This effect is potentially dependent on released bacterial ATP given that no differences in bacterial counts were observed (see Fig. 4D and 4E, Figure 4-figure supplement 1D-E).

In summary, ATP released by bacteria suppresses abdominal inflammatory responses and impaired survival of mice in a model of abdominal sepsis.

### Establishing ATP-loaded OMV as a model system to assess the systemic relevance of bacterial ATP

ATP is rapidly metabolized in the extracellular space and therefore, the mode of action of released bacterial ATP is limited to the immediate cellular vicinity (Junger, 2011). However, the outcome of sepsis is not only dependent on local but also on systemic responses to microorganisms. Therefore, we hypothesized that bacterial ATP has systemic effects as protected cargo in OMV (Alvarez et al., 2017). OMV are small (20-300nm) spherical particles that are released by both gram^neg^ and gram^pos^ bacteria (Schwechheimer & Kuehn, 2015). In gram^neg^ bacteria, they bulge off the outer membrane and disseminate throughout the body (Jang et al., 2015). They are equipped with typical bacterial surface features lacking the machinery for self-reproduction, and contain DNA, proteins, and metabolites (Baeza & Mercade, 2021; Bitto et al., 2017; Kulp & Kuehn, 2010; E.-Y. Lee et al., 2007). Therefore, OMV are suited as a systemic delivery system for bacterial ATP. Indeed, recently, ATP has been detected in OMV derived from pathogenic *Neisseria gonorrhoeae*, *Pseudomonas aeruginosa* PAO1 and *Acinetobacter baumannii* AB41 (Pérez-Cruz et al., 2015).

To assess the potential of OMV as ATP carriers, we compared the OMV production from several hypervesiculation *E. coli* mutants (Δ*mlaE*, Δ*mlaA*, Δ*rfaD*, Δ*degP*, Δ*rodZ*, Δ*nlpI*, Δ*tolB*) (Figure 5A) (McBroom et al., 2006). The Δ*nlpI* and Δ*tolB* strains showed a 20- and 30-fold increase of OMVs when compared with the PS (Figure 5B). We then assessed ATP release and growth over time from Δ*nlpI*, Δ*tolB* and the PS, to identify their individual peak of ATP release and isolated OMV at their individual peak of ATP release and after 24h (Figure 5-figure supplement 1A). ATP was detected in OMV from all assessed strains at the individual peak of ATP release but only in minimal detectable levels after 24h (Figure 5C). The Δ*tolB* OMV isolated after 24h were then used as ATP-depleted vehicles. After density gradient ultracentrifugation, most Δ*tolB* OMV were either in fraction 3a or 3b and therefore of similar density (Figure 5D, Figure 5-figure supplement 1B). They were equipped with outer membrane ompF but not cytoplasmic ftsZ (Figure 5D), indicating that they are outer membrane derived. In order to generate OMV with known and constant ATP concentrations, we used electroporation to load OMV with ATP while empty mock-electroporated OMV (Δ*tolB* OMV harvested from 24h culture) served as ATP-depleted controls (Fu et al., 2020; Lennaárd et al., 2021). Before and after electroporation, the OMV size distribution was assessed by nanoparticle tracking analysis and morphology by electron microscopy (Figure 5E, Figure 5-figure supplement 1C). OMV were loaded with ATP (Figure 5F) and over time, the amount of ATP in OMV decreased, especially at physiologic (37°C) temperature (Figure 5G) as opposed to 4°C (Figure 5-figure supplement 1D).

**Figure 5.**
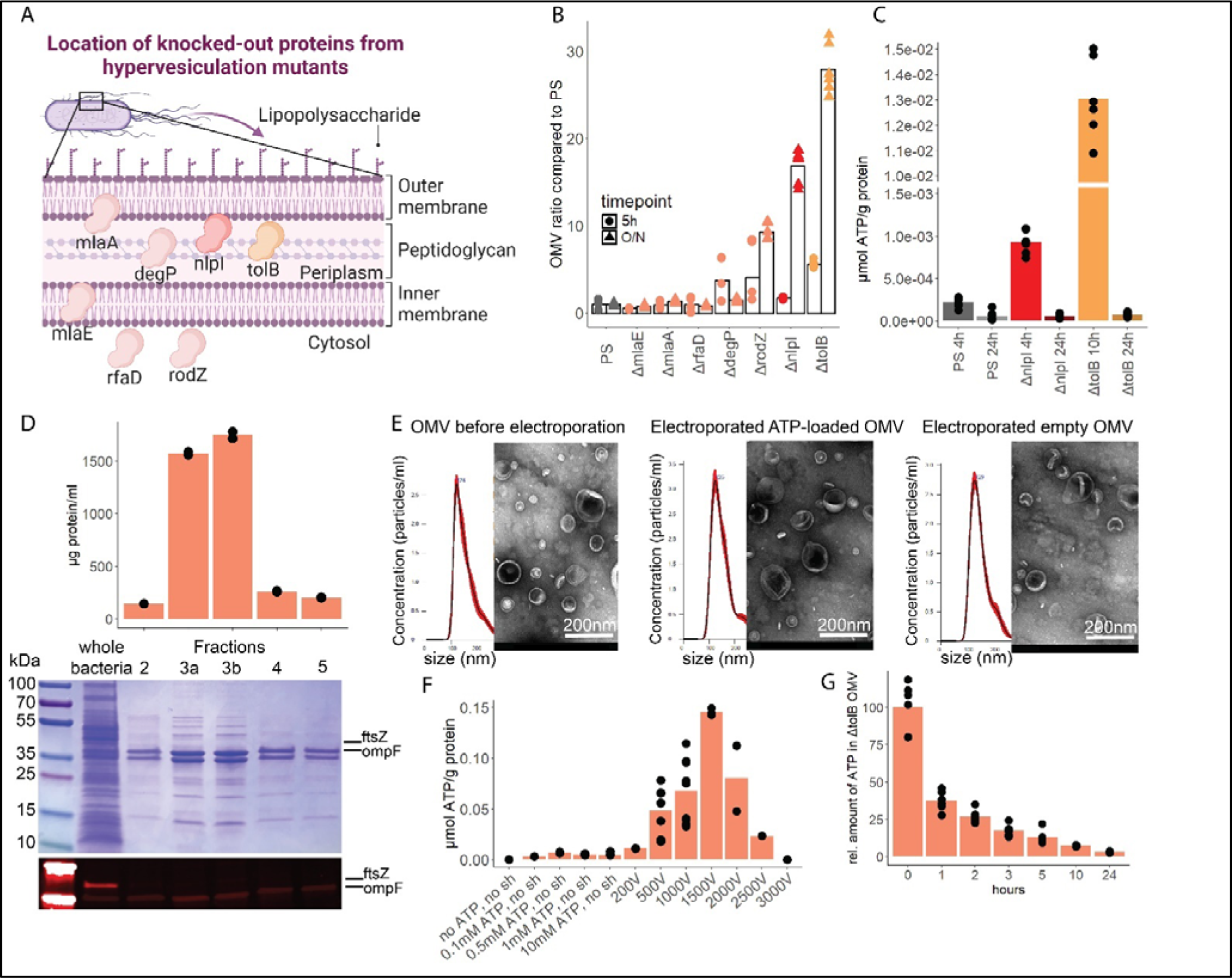
OMV contain ATP and can be exploited as a model to assess the systemic relevance of bacterial ATP. **(A)** Illustration depicting the location of assessed proteins that lead to a hypervesiculation phenotype if knocked-out in the gram^neg^ bacterium *E. coli*. **(B)** Relative amount of OMV (µg protein/ml) compared to the PS isolated from growth cultures of the assessed hypervesiculation mutants after five hours (exponential growth) and O/N (stationary phase). n = 2 measurements of N = 3 independent bacteria cultures. Means and individual values are shown. **(C)** Absolute quantification of ATP in OMV isolated from growth cultures of the PS, Δ*nlpI* and Δ*tolB* at their individual peak of ATP release and after 24 hours. n = 2 measurements of N = 3 independent bacteria cultures. Means and individual values are shown. **(D)** Amount of protein (BCA assay) detected in different fractions after density gradient ultracentrifugation. n = 2 measurements of the different fractions. 20µl of *E. coli* growth culture and 20µl of each fraction were then characterized by Coomassie blue staining and specific detection of outer membrane ompF and cytoplasmic ftsZ. **(E)** Characterization of OMV by nanoparticle tracking analysis (n = 5 measurements per sample) and electron microscopy (representative image) before and after electroporation. **(F)** Absolute quantification of ATP in OMV, which were loaded using different strategies. Column 2-5: different concentrations of ATP incubated for one hour at 37°C (passive filling). Column 6-12: different voltages with fixed settings for Resistance (100Ω) and Capacitance (50µF). N = 2-9 independent experiments. Means and standard deviations are shown. **(G)** Relative quantification of ATP in OMV over 24 hours at 37°C after electroporation (0h = 100%). N = 2 measurements of N = 3 independent experiments. Means and individual values are shown.

In summary, OMV contain ATP and release ATP at physiological temperatures. To use OMV as an ATP delivery system, empty OMV were loaded using electroporation.

### OMV-derived bacterial ATP induces degranulation processes in neutrophils after lysosomal uptake

OMV are potent inducers of inflammation and sepsis (Park et al., 2010, 2013), which travel throughout the body and are taken up by a variety of cells (Bittel et al., 2021; Kim et al., 2013; J. Lee et al., 2018). To tests the hypothesis that ATP within OMV mediates systemic effects of invasive bacteria, we injected ATP-loaded and empty OMV i.a. and investigated the resulting inflammation one hour after injection (Figure 6A).

**Figure 6.**
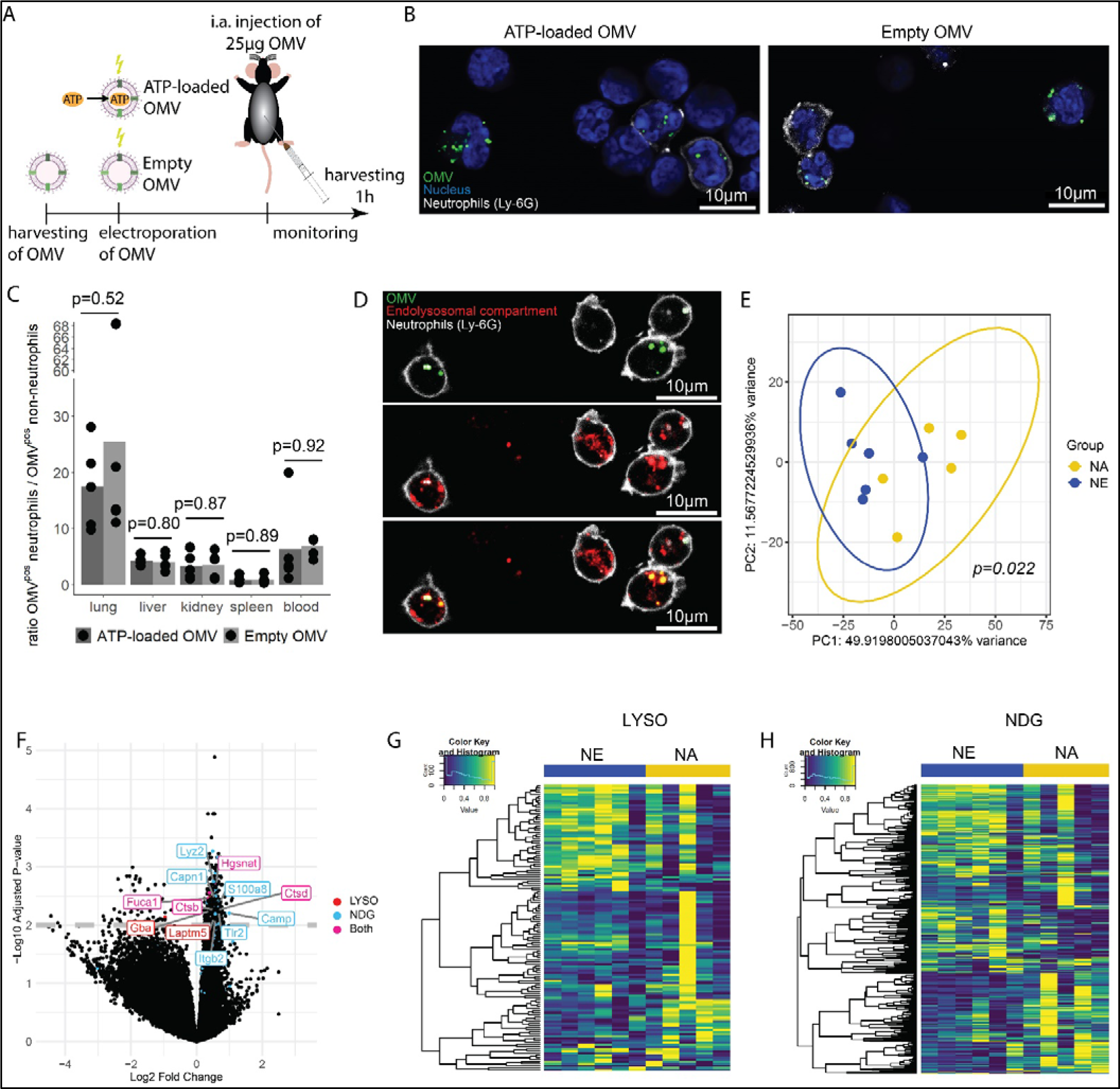
Bacterial ATP within OMV upregulates lysosome-related pathways and degranulation processes in neutrophils. **(A)** Experimental approach to determine the systemic role of bacterial ATP *in vivo*, i.a. injecting ATP-loaded or empty OMV. **(B)** Representative microscopic images of cells from the abdominal cavity one hour after i.a. injection of either ATP-loaded or empty OMV. OMV: DiI, Nucleus: DAPI, Neutrophils: Ly-6G-FITC. **(C)** Cells from remote organs were isolated one hour after i.a. injection of either ATP-loaded or empty OMV. OMV were mainly taken up by neutrophils (except in the spleen, ratio ≈ 1). T-test with Benjamini-Hochberg correction, n = 5 animals per group of N = 2 independent experiments. Means and individual values are shown. **(D)** Representative microscopic image of pulmonary neutrophils one hour after i.a. injection of either ATP-loaded or empty OMV. OMV co-localize with the endolysosomal compartment. OMV: DiI, Endolysosomal system: Deep Red LysoTracker, Neutrophils: Ly-6G-FITC. **(E)** Pulmonary neutrophils were isolated one hour after i.a. injection of ATPγs-loaded or empty OMV, bead-sorted and RNA sequencing was done. Principal component analysis shows significantly different clustering between neutrophils that took up ATPγs-loaded (NA) or empty OMV (NE). PERMANOVA, n = 6 animals in the NE group, n = 5 animals in the NA group. Ellipses represent 95% confidence level. **(F)** Volcano plot of RNA sequencing results shows an upregulation of genes mainly in the NA group. Genes classified in either lysosome (LYSO) or neutrophil degranulation pathways (NDG) or both, which were mentioned in the text, were highlighted. **(G)** Heatmap of the lysosome pathway (LYSO) showing the gene expression per sample. **(H)** Heatmap of the neutrophil degranulation pathway (NDG) showing the gene expression per sample.

In the abdominal fluid, uptake of DiI-stained OMV by leucocytes was independent of ATP cargo (Figure 6B, Figure 6-figure supplement 1) and all major cell populations with phagocytic ability (LPM, SPM, neutrophils) were highly positive for OMV (Figure 6-figure supplement 2A-B). LPM dramatically decreased, whereas neutrophils increased in response to OMV when compared to sham controls (Figure 6-figure supplement 2C).

Within one hour, the DiI-stained OMV were distributed throughout the body and mainly taken up by neutrophils, with remarkable differences between organs (Figure 6-figure supplement 3A-C). The highest ratio of OMV^pos^ neutrophils compared to all other OMV^pos^ cells was observed in the lung in an ATP-independent manner (Figure 6C). Intracellular fate of engulfed OMV is still a matter of discussion but one possibility is the degradation in the endolysosomal compartment (Bielaszewska et al., 2017; Juodeikis & Carding, 2022). We therefore injected DiI-stained OMV i.a. to assess to which membrane compartment OMV co-localize. A strong co-localization with the endolysosomal compartment of pulmonary neutrophils was identified (Figure 6D).

Next, we aimed to determine what pathways OMV-derived bacterial ATP initiates in neutrophils after uptake into the endolysosomal compartment. During infection and sepsis, neutrophils are highly abundant in the lungs and most OMV were engulfed by neutrophils (see Figure 6C and Figure 6-figure supplement 3C). Because of its anatomically central position in the blood circulation and extensive vasculature, the lung is often affected in sepsis and a significant cause of mortality (Wang et al., 2024). We therefore isolated again pulmonary neutrophils (Figure 6-figure supplement 4) for RNA sequencing. Uptake of ATP-loaded OMV resulted in a significantly different transcriptional activity compared to empty OMV (Figure 6E): The strongest differences were found in lysosomal (LYSO) and neutrophil degranulation pathways (NDG) (Figure 6F, Figure 6-figure supplement 5). In the lysosomal pathway, transcripts for proteins involved in autophagosome-lysosome fusion (Gabarap), endosome to lysosome transport (Hgsnat, Laptm5), or enzymatic cleavage (Capn1, Fuca1, Ctsb, Ctsd, Gba) were significantly higher expressed in response to ATP-loaded OMV when compared to empty OMV (Figure 6F-G). Furthermore, neutrophil degranulation was significantly increased in response to ATP-loaded OMV as indicated by increased expression of gene transcripts resulting in antibacterial activity (Lyz2, S100a8, Camp) or NETosis (Tlr2, Itgb2) (Figure 6F and H).

In summary, OMV are phagocytosed locally by major cell populations with phagocytic ability. Remotely, OMV were mainly taken up by neutrophils, where they co-localize with the endolysosomal compartment. Delivery of ATP by OMV resulted in an upregulation of lysosomal activation and neutrophil degranulation.

## Discussion

In this study, we have demonstrated that ATP release is dependent on the respiratory chain located at the inner bacterial membrane. ATP synthase is most likely dominant compared to cytochrome *bo_3_* oxidase because of its non-redundant function for ATP generation and bacterial growth. Mutations in one of the cytochrome *bo_3_* oxidase subunits cyoA, cyoC or cyoD could be compensated by another subunit. This is supported by lower ATP release in the cyoB mutant, which harbors all three prosthetic groups of the cytochrome *bo_3_* oxidase and is therefore indispensable for proper function (Tsubaki et al., 2000). Alternatively, deficiency in the *bo_3_*-type oxidase could be partially compensated by *bd*-type oxidases (Borisov et al., 2011; Grauel et al., 2021).

Instability of the outer bacterial membrane as we have seen for the Δ*ompF* mutant (Choi & Lee, 2019) is associated with bacterial death and ATP release. It remains unclear whether loss of ATP results directly in bacterial death or bacterial death is the direct result of impaired outer bacterial membrane stability and ATP release is secondary to bacterial death. However, since the inner membrane and ATP generation remains intact and the periplasmic space is considered generally devoid of ATP, the latter seems more likely (Kulp & Kuehn, 2010). Previous studies used Sytox/PI staining and microscopy to identify bacterial death and did not identify bacterial death as a relevant source of extracellular ATP during exponential growth (Hironaka et al., 2013). Conversely, we used flow cytometry, a quantitatively more sensitive method and our data demonstrated that impaired membrane integrity and bacterial death are critical for ATP release during growth. Despite low rates of bacterial death during exponential growth (Koch, 1959), and a low ratio of dead to live bacteria, the high gradient between intra- and extracellular ATP (approximately 100 - 1000x higher) might be sufficient to explain the amount of released ATP measured (Leduc & Van Heijenoort, 1980; Mempin et al., 2013).

In this study, we used the above-mentioned outer membrane porin mutants to study impaired membrane integrity and we used hypervesiculation mutants to study the effects of OMV-derived bacterial ATP. It is important to note that these impaired membrane integrity and hypervesiculation phenotypes are not clearly distinct but rather overlapping. However, outer membrane porin mutants are not known as high OMV producers. Thus, measured ATP from outer membrane porin mutants was most likely directly released and not OMV-derived ATP. To assess ATP within OMV, we switched to typical hypervesiculation mutants.

ATP was detected within OMV and the concentration of ATP within OMV is sufficient to activate P2-type receptors in lysosomes (Araujo et al., 2020; Junger, 2011). Furthermore, OMV are released in response to environmental stresses like infection and sepsis (Kulp & Kuehn, 2010; Orench-Rivera & Kuehn, 2016). ATP is likely to activate P2 receptors, which have been shown to be expressed in lysosomes, such as P2X1, P2X3 or P2X4 (Qureshi et al., 2007; Robinson & Murrell-Lagnado, 2013). OMV can therefore be considered as non-living bacteria-resembling highly inflammatory vesicles (Jang et al., 2015; Kulp & Kuehn, 2010) that effectively distribute ATP throughout the body to activate cells, such as neutrophils, initiating systemic inflammatory responses. It has already been shown that OMV are engulfed by neutrophils (Sartorio et al., 2021), however, the finding that they actually have a functional effect is a novelty. Activation of neutrophils in response to ATP-loaded OMV resulted in upregulation of inflammatory, lysosomal, and neutrophil degranulation pathways as well as an upregulation of the apoptosis pathway and genes involved in NETosis by a hitherto unknown mechanism. These pathways explain in part the low neutrophil counts we observed in mice. Such remote degranulation in response to OMV is unlikely to control bacteria at the source of sepsis but rather harmful to host tissues increasing sepsis severity (Eichelberger & Goldman, 2020).

The study has several limitations: Bacterial ATP release is strain-specific (Figure 1) (Mempin et al., 2013). As we focused on the laboratory BW25113 *E. coli* strain, it remains to be elucidated if impaired outer membrane integrity and bacterial death are also of such importance in other gram^neg^ or also gram^pos^ bacteria. The approach to load OMV with ATP is rather artificial; it is, however, the only way to assure that ATP-loaded and empty OMV only differ in their ATP cargo. OMV have surface antigens and contain DNA, proteins and other metabolites (Kulp & Kuehn, 2010), which are known to elicit inflammation as well. This was controlled using the same OMV as baseline vehicles and including electroporation also for the controls.

Given the magnitude of the difference in concentrations we detected between directly released and OMV-derived ATP, the role of OMV-derived ATP was not assessed locally, and we focused on its remote effects. However, the signaling mechanisms between directly released ATP (extracellular) and OMV-delivered ATP (endolysosomal compartment) are different. Therefore, the local role of OMV-derived ATP should be explored in future studies, including potential differences in transcription or cytokine profiles.

Other open questions need to be addressed in future projects: It remains to be determined how OMV are physiologically loaded with ATP. Despite the lack of a transporter, ATP may leak across the inner membrane into the periplasmic space. Such leakage may be similar to baseline leakage in eukaryotic cells (Yegutkin et al., 2006). Different types of OMV have been described in recent years, for example outer-inner membrane vesicles (O-IMV) or explosive OMV (Juodeikis & Carding, 2022; Takaki et al., 2020; Toyofuku et al., 2019; Turnbull et al., 2016; Turner et al., 2015), which are composed of outer membrane, inner membrane and cytoplasmic content. The mechanisms, how these types of OMV are generated, could explain that ATP is found within them. However, the Δ*tolB* mutant produces, unlike other tol mutants, only very few O-IMV (0.1%-2% of all OMV) (Pérez-Cruz et al., 2013; Reimer et al., 2021; Takaki et al., 2020) and our Western blot analysis suggests that our OMV are primarily outer membrane derived. Future studies may address the ATP cargo of the different OMV subgroups. Another open question is generally the physiological role of OMV-delivered ATP. We detected concentrations around 1-10 nmol/g protein, which is in accordance with Pérez-Cruz et al. (Pérez-Cruz et al., 2015). Interestingly, Alvarez et al. reported 1000x higher concentration (∼13nmol/µg protein). This difference might be explained by the techniques used. Considering our chosen voltage of 1100V and the ATP decrease in OMV over time, we reach similar concentration as measured in our isolated OMV. It is therefore reasonable to assume that our shown effects are physiological, which has, however, to be confirmed by other studies.

OMV are promising diagnostic molecular biomarkers in gram^neg^ sepsis (Michel & Gaborski, 2022) and it would be of relevance to compare OMV isolated from septic and control patients to assess possible differences in ATP cargo. Furthermore, it remains to be elucidated, which mechanisms lead to low local neutrophil counts. Since ATP is involved in cell death (Proietti et al., 2019b) as well as in chemotaxis (Junger, 2011), increased cell death, impaired infiltration or a combination of both is possible.

This study reveals that ATP is released by bacteria during growth because of impaired membrane integrity and bacterial death. ATP is also being released via OMV and therefore acts locally (direct release) and systemically (via OMV). Bacterial ATP reduces neutrophil counts, activates the endolysosomal system and upregulates neutrophil degranulation, which together increase the severity of abdominal infection and early sepsis. These findings have the potential to lead to the development of novel treatments for abdominal sepsis, e.g. by *in vitro* generated OMV that modulate neutrophil function via delivery of inhibitors to intracellular purinergic receptors during sepsis.

## Materials and Methods

### Human data

From five patients that underwent revision laparotomy because of abdominal sepsis, swabs were taken from abdominal fluid, streaked on LB agar plates (15g Agar, 5g Bacto Yeast Extract, 10g Bacto Tryptone, 5g NaCl in 1L ddH_2_O; key resources table) and cultivated an-/aerobically for 48h. The human experimental protocol was approved by the Cantonal Ethics Commission Bern, Switzerland (ethical approval 2017-00573, NCT03554148). Written informed consent was obtained from all patients and the study has been performed in accordance with the Declaration of Helsinki as well as the CONSORT statement.

### Mouse handling

Specific-pathogen-free C57Bl/6JRccHsd mice (key resources table) were purchased at the age of 8 weeks from Inotiv (earlier Envigo, the Netherlands) and were housed in ventilated cages in the Central Animal Facility, University of Bern, Switzerland. All experiments were performed in the morning, mice were supplied with a 12-hour light/dark cycle at 22°C and fed ad libitum with chow and water. To minimize cage effects, we mixed mice over several cages and therefore only used female mice. All animal procedures were carried out in accordance with the Swiss guidelines for the care and use of laboratory animals as well as in accordance with the ARRIVE guidelines and were approved by the Animal Care Committee of the Canton of Bern (Switzerland) under the following number: BE41/2022.

### Cecal ligation and puncture (CLP) sepsis model

To isolate bacteria from mice with abdominal sepsis, cecal ligation and puncture was performed as described elsewhere with some minor modifications (Rittirsch et al., 2009). In brief, mice were anesthetized s.c. injecting (3µl/g body weight) a mixture of fentanyl (0.05mg/ml), dormicum (5mg/ml) and medetor (1mg/ml) and were then shaved and disinfected with Betadine. Mid-line laparotomy was performed (approx. 1 cm) and the cecum was exposed. The proximal 1/3 of the cecum was ligated with Vicryl 4-0 (Ethicon, cat# V1224H) and perforated with a 23 G needle. The cecum was returned to the abdominal cavity and the laparotomy was sutured continuously in two layers with prolene 6-0 (Ethicon, cat# MPP8697H). At the end, the antidote (naloxone (0.1mg/ml), revertor (5mg/ml), temgesic (0.3mg/ml)) was s.c. injected (9µl/g body weight). A semi-quantitative score sheet was used to predict animal postoperative well-being. Mice were evaluated every four hours according to the following criteria: appearance, level of consciousness, activity, response to stimulus, eye shape, respiratory rate and respiratory quality and analgesia was applied if necessary. After 10 hours, abdominal fluid was collected, spread on LB agar plates, and cultivated an-/aerobically for 48h.

### Whole 16S-rRNA sanger sequencing

Twenty-five colonies cultivated from abdominal fluid of patients and mice each described above were randomly picked and collected in separate sterile Eppendorf tubes and resuspended in 1ml of sterile PBS. Each sample was centrifuged for 5 min. at 20’000g and washed once with 1ml sterile PBS. The pellet was resuspended in 20µl of sterile PBS and samples were incubated 5min. at 100°C. 1µl was used for PCR using GoTaq G2 Green Master Mix (key resources table) and the following primers (key resources table) at a final concentration of 0.2µM:

-fD1: 5’-AGA-GTT-TGA-TCC-TGG-CTC-AG-3’

-fD2: 5’-AGA-GTT-TGA-TCA-TGG-CTC-AG-3’

-rP1: 5’-ACG-GTT-ACC-TTG-TTA-CGA-CTT-3’

PCR conditions were as follows: Initial 5min. at 94°C for denaturation, followed by 35 cycles of 1min. denaturation at 94°C, 1min. annealing at 43°C, and 2min. extension at 72°C, with a final extension for 10min at 72°C. This resulted in a PCR product of optimally ∼1400 - 1500bp. 20µl PCR product were run on 2% agarose gel for 90min., cut out and purified using Qiaquick Gel Extraction Kit (key resources table). The amplicon concentration was measured using nanodrop (Thermo Fisher Scientific) and sanger sequencing was done by microsynth.

### Quantification of released bacterial ATP

Bacteria were aerobically grown in 20ml LB medium (5g Bacto Yeast Extract, 10g Bacto Tryptone, 5g NaCl in 1L ddH_2_O; key resources table) overnight (O/N) (16h / 37°C / 200rpm) and 0.25ml of O/N culture were diluted in 100ml fresh LB medium. Up to 24 hours, 0.25ml bacteria culture were taken at several time points for growth assessment (OD_600_) and 1ml was taken for ATP quantification. OD_600_ was measured using a Tecan Spark spectrophotometer. The 1ml sample for ATP quantification was centrifuged for 5min. at 16’000g, the supernatant was filtered through a 0.2µm syringe filter and stored at -80°C. ATP was quantified using a luciferin-luciferase-based assay according to the manufacturer’s protocol (ATP Kit SL, key resources table) and bioluminescence was measured using a Tecan Spark spectrophotometer. In Figure 3, LB medium was supplemented with 1mM calcium or 0.5mM EDTA for the controls.

### Absolute quantification of bacteria and assessment of viability

Growth culture was set up as described above and after 4 hours (ATP peak), samples were taken to quantify bacteria and assess viability. Bacteria were diluted in PBS and stained using the Cell Viability Kit with BD Liquid Counting Beads (key resources table) according to the manufacturer’s protocol. In brief, 100µl of diluted bacterial growth culture was stained with 1µl TO dye, 1µl PI dye and 10µl of counting beads were added. The sample was acquired on a CytoFlex S (Beckman Coulter), setting the thresholds for the TO- and the PI-channel to 1000 and analysis was done with FlowJo software (key resources table). Bacterial biomass was calculated to bacteria/ml culture according to the manufacturer’s protocol.

### Transformation of *E. coli* PS

The Keio collection (key resources table) *E. coli* PS was aerobically grown in LB medium O/N (16h / 37°C / 200rpm). 0.5ml O/N culture was diluted in 35ml fresh LB medium and incubated until OD_600_ was 0.35-0.45. Cultures were chilled in ice-water and washed two times with ice-cold 20ml ultra-pure water (Thermo Fisher Scientific, cat# 10977035) (20min. / 4°C / 3200g). The supernatant was carefully decanted, and the bacterial pellet was gently resuspended by pipetting (no vortex). After the final wash, the pellet was resuspended in 240µl ice-cold ultra-pure water and kept on ice until electroporation. 80µl of ice-cold bacterial cells were mixed with 1µl plasmid (pBAD28 == pEMPTY or pHND10 == pAPY, 100-200ng, key resources table) in chilled 0.1cm cuvettes (Bio-Rad, cat# 1652089) and immediately electroporated (Voltage = 1.8kV, Capacitance = 25µF, Resistance = 200Ω). Directly after electroporation, 1ml of warm LB medium was added to the cuvette, bacteria were transferred to a tube containing 10ml pre-warmed LB medium and cultivated for 1h at 37°C. Then, bacteria were dispersed on LB agar plates supplemented with ampicillin (100 µg/ml) and incubated at 37°C. The next day, three colonies were picked, streaked on a new ampicillin supplemented LB agar plate, and incubated for 24h. The following day, two cryostocks were made from single colonies in LB medium supplemented with 20% glycerol.

### Bacteria injection sepsis model

The Keio collection (key resources table) *E. coli* PS transformed with either pEMPTY or pAPY was aerobically grown O/N (16h / 37°C / 200rpm) in 20ml LB medium supplemented with ampicillin (100 µg/ml). PS+pEMPTY and PS+pAPY were washed with PBS (20min. / 22°C / 3200g) and resuspended in 20ml fresh LB medium supplemented with ampicillin (100 µg/ml). To induce the apyrase, L-arabinose (0.03%, Sigma-Aldrich, cat# A3256-25G) was added. After three hours, bacteria were washed, resuspended in PBS supplemented with L-arabinose (0.03%) and 2 x 10^9^ colony forming units were injected i.a. ATP in bacteria cultures and in the abdominal fluid was assessed using the same assay as described above (ATP Kit SL, key resources table). Mice were evaluated for post-operative well-being every four hours as described above. After four and eight hours and when the score reached specific criteria (for the survival experiment), animals were sacrificed using pentobarbital (150mg/kg bodyweight) followed by organ collection.

### Collection of abdominal fluid, blood and organs

After pentobarbital injection, the mice were placed on a surgical tray, fixed, the abdomen was shaved, the abdominal skin was disinfected with betadine and the skin (but not peritoneum) was cut. Abdominal cells were isolated as described elsewhere with some modifications (Ray & Dittel, 2010). In brief, a 22 Gauge peripheral IV catheter was inserted into the abdominal cavity through the peritoneum. The abdominal cavity was flushed two times with 5ml MACS buffer (PBS supplemented with 3% FBS (Gibco, cat# 10500-064), 2% HEPES (Sigma-Aldrich, cat# H0887-100ml) and 2mM EDTA (Sigma-Aldrich, cat# E5134-500G)). The first 5ml were used to flush the upper abdomen under pressure, and the second 5ml, to flush the lower abdomen under pressure. Part of the aspirated fluid was used for aerobic plating, if needed, and the rest was centrifuged for 5min at 700g to pellet the abdominal cells for various downstream applications. To collect blood, the peritoneum was opened. 300µl blood was collected from the inferior vena cava using a 22 Gauge peripheral IV catheter and a 1ml syringe, which was prefilled with 30µl 2mM EDTA. Before organ collection (lungs, liver, kidney, and spleen), the mouse was flushed with 5ml PBS. Organs were excised and used for downstream processing.

### Digestion of organs and preparation of single cell suspension

Harvested organs were digested (key resources table; kidney&lungs: 1mg/ml Col I + 1mg/ml Col IV + 1mg/ml Col D + 0.1mg/ml DNase I in DMEM (Gibco, cat# 31966-047) + 3% FBS for 30 min.; liver: 1mg/ml Col IV + 0.1mg/ml DNase I in DMEM + 3% FBS for 20 min.) at 37°C with a spinning magnet or gently pushed through a 100µm mesh (spleen). After washing with MACS buffer (5min. / 4°C / 700g), erythrocytes were lysed using self-made RBC buffer (90g NH_4_Cl, 10g KHCO_3_, 370mg EDTA in 1L ddH2O for 10x stock). Cells were washed again and stained as described below.

### Staining of cells and flow cytometry

First, viability dye (key resources table) and Fc-block (key resources table) were diluted in PBS and cells were incubated for 20min. at 4°C in the dark. Cells were washed with MACS buffer and surface staining was done with the listed antibody cocktail: (key resources table: Ly-6G FITC, Ly-6C PerCP-Cy5.5 or Ly-6C PE-Cy7, CD11b APC, CD206 AF700, CD11c APC-eFluor780, CD45 efluor450 or CD45 APC-Cy7, CD19 Super Bright 600, CD3 BV605, NK1.1 BV605, CCR2 BV650, I-A/I-E BV711, CX3CR1 BV785, Siglec F PE, FcεR1α PE/Dazzle 594, CD115 PE-Cy7, F4/80 BUV395) for 20min. at 4°C in the dark. Cells were washed again and resuspended in MACS buffer for acquisition on an LSR-Fortessa (BD Biosciences). Analysis was done with FlowJo software (key resources table) and OMIQ web-based analysis platform (https://www.omiq.ai/). To preserve the global structure of abdominal cell populations, uniform manifold approximation and projection (UMAP) was used as dimensionality reduction technique and cell populations were defined using FlowSOM clustering algorithm (Van Gassen et al., 2015).

### Absolute quantification of bacteria by plating

To count bacteria in isolated abdominal fluid, blood or growth cultures, serial dilutions were done (1 to 1:100’000) and 50µl of each dilution was streaked on LB agar plates and aerobically incubated. Plates, which had between 20-200 colony forming units were used for quantification.

### Collection of OMV and ATP measurement of OMV

*E. coli* PS or hypervesiculation mutants from the Keio collection were grown in in LB medium for 5h, O/N (16h) or 24h at 37°C and 200rpm. Bacteria cultures were then centrifuged (20min. / 4°C / 3200g) to pellet bacteria. The supernatant was filtered through a 0.45µm PES filter (key resources table) and ultracentrifuged (1.5h / 4°C / 150’000g) to pellet OMV. When ATP within OMV was measured, OMV were washed in PBS and directly stored at -80°C. ATP was quantified using a luciferin-luciferase-based assay according to the manufacturer’s protocol (Microbial ATP Kit HS, key resources table) and bioluminescence was measured using Tecan Spark spectrophotometer.

If OMV were used for characterization or i.a. injection, first ultracentrifugation was followed by a density gradient ultracentrifugation using OptiPrep (Iodixanol, Stemcell technologies, cat# 07820). OMV were resuspended in 50% OptiPrep and OptiPrep gradient (10%, 20%, 30%, 40%, 45%, 50%) was made in underlay-technique starting with the 10% layer. Samples were ultracentrifuged (16h / 4°C / 150’000g) and 6 fractions (fractions 1, 2, 3a, 3b, 4, and 5) were defined. Fractions were washed separately with PBS (1.5h / 4°C / 150’000g) and the amount of OMV was determined measuring protein concentration using BCA assay according to the manufacturer’s protocol (Thermo Fisher Scientific, cat# 23227). For experiments, only OMV from fraction 3 were used, which were washed with PBS (1.5h / 4°C / 150’000g), resuspended in PBS and stored at -80°C until further processing.

### Electroporation (EP) and staining of OMV

If electroporation and staining of OMV was performed, OMV were thawed, pelleted (1.5h / 4°C / 150’000g), the pellet was resuspended in 720µl EP-buffer (500mM Sucrose and 1ml Glycerol in 10ml ultra-pure water) and kept on ice. This suspension was mixed with either 80µl EP-buffer (control) or 80µl 10mM ATP (or ATPγs for RNA sequencing experiment) (key resources table), filled in chilled 0.4cm EP cuvettes (Bio-Rad, cat# 1652088) and immediately electroporated (Voltage = 1100V, Capacitance = 50µF, Resistance = 100Ω). After EP, the OMV were 1:1 diluted in warm PBS and kept at 37°C for 20min. DiI (key resources table) was added 1:100 to the sample during the incubation time. OMV suspension was then washed with PBS (1.5h / 4°C / 150’000g).

### OMV injection sepsis model

After the final wash in PBS (see above), OMV were resuspended in NaCl 0.9% and filtered through a 0.45µm centrifuge filter tube (Sigma-Aldrich, cat# CLS8162-96EA). The final OMV suspension was quantified using a BCA assay. 25µg of either ATP-loaded OMV or empty OMV were i.a. injected and mice were evaluated as described above. After one hour, the animal was sacrificed using pentobarbital followed by organ collection. Tissue digestion and flow cytometry was done as described above.

### Assessment of ATP release by OMV

OMV were electroporated as described above. After washing in PBS (1.5h / 4°C / 150’000g), the OMV pellet was resuspended in 6ml warm PBS and incubated at 37°C. At baseline, after 1, 3, 5, 10 and 24 hours, 1ml was taken and washed with PBS (1.5h / 4°C, 150’000g). The OMV pellet was then resuspended in 200µl PBS to assess protein concentration using a BCA assay, and ATP in OMV was quantified using a luciferin-luciferase-based assay according to the manufacturer’s protocol (Intracellular ATP Kit HS, key resources table). Bioluminescence was measured using a Tecan Spark spectrophotometer.

### SDS-PAGE, protein staining and Western blot

20µl of diluted *E. coli* culture and 20µl of the different OMV fractions (except fraction 1, since no protein could be detected) were diluted 1:1 with Laemmli/βME (Laemmli buffer solution containing 5% β-mercaptoethanol). The mixture was heated at 100°C for 5min., shortly centrifuged at 13’000rpm and loaded on Mini-PROTEAN TGX Gels (Bio-Rad, cat# 4561094). Bio-Rad marker (6µl) was added to one well and the gel was run at 100V for 1.5h. The gel was then either directly stained with Coomassie blue or transferred to a membrane.

Coomassie blue staining was done as follows: The gel was washed in ddH_2_O and the staining solution (Coomassie blue 0.1%, 40% ethanol, 10% acetic acid) was heated for 15s in the microwave. Warm staining solution was added to the gel and gently shaken for 15min. Then, staining solution was removed, and the gel was washed with ddH_2_O. Destaining solution (10% ethanol, 7.5% acetic acid) was added for 1h, exchanged and left overnight.

To transfer the proteins to a membrane, iBlot2 (Invitrogen) was used. Membranes were then blocked with 4% milk/PBS for 1h. FtsZ-antibody (1:200, key resources table) and ompF-antibody (1:500, key resources table) were added and incubated overnight at 4°C. The next day, membranes were washed with PBS tween (PBST, 0.05%) three times for 5min. Secondary fluorescent antibody was then added in milk (1:10’000, key resources table) and membrane was incubated for 1h. The membrane was washed again with PBST three times for 5min. and then scanned using a Licor Odyssey.

### Nanoparticle tracking analysis (NTA)

Size distribution of OMV was analysed using the NanoSight NS300 Instrument (Malvern Panalytical,405 nm laser) according to the manufacturer’s protocols. OMV were resuspended in PBS and serial dilutions (1 to 1:100’000) were used to find suitable concentrations. Each experimental sample was analyzed 5 times. PBS was used to flush the system between the samples and to assess background. For each sample, the relative amount of OMV and the OMV size was recorded, which resulted in a size distribution curve. NTA 2.3 software was used to analyze the data and the following script was used for acquisition (Gheinani et al., 2018): SETTEMP 25; CAMERAON; CAMERAGAIN 12; CAMERALEVEL 11; REPEATSTART; SYRINGLOAD 100; DELAY 10; SYRINGSTOP; DELAY 15; CAPTURE 60; DELAY 1; **REPEAT 4**; SETTEMP OFF; PROCESSINGLESETTING; EXPORTRESULTS.

### Electron microscopy negative staining

For imaging of negatively stained samples, 5µl of OMV suspension were adsorbed on glow discharged carbon coated 400 mesh copper grids (Plano) for 1-5min. After washing the grids 3 times by dipping in ultra-pure water, the grids were stained with 2% uranyl acetate solution (Electron Microscopy Science) in water for 45 s. The excess fluid was removed by gently pushing the grids sideways onto filter paper. The grids were then examined with a FEI Tecnai Spirit transmission electron microscope at 80kV, which was equipped with a Veleta TEM CCD camera (Olympus).

### Immunofluorescent microscopy

One hour after i.a. injection of DiI-stained OMV, lungs were digested as described above. Single cell suspension was either fixed and imaged or live cells were imaged. For the fixed approach, single cell suspension was immunolabeled with FITC-tagged anti-Ly-6G (key resources table) at 1:100 dilution for 20 minutes at 4°C in the dark to distinguish neutrophils. Cells were washed once with PBS and applied onto glass slides using Cytospin (Thermo Shandon) for 5 minutes (1800 rpm). Slides were air dried for 2 minutes, washed with PBS and fixed using 4% paraformaldehyde solution for 10 min. Cell nuclei were stained with DAPI (key resources table) at 1:5000 concentration in IF buffer (0.25% BSA, 0.1% Triton X-100 in PBS) for 1.5 hours at room temperature in the dark. After washing 3×5 min. with IF buffer, the slides were covered and sealed with nail polish.

If live cells were imaged, single cell suspension was concurrently immunolabeled and stained with Hoechst 33342 (key resources table) at 1:1000 dilution for cell nuclei, FITC-tagged anti-Ly-6G (key resources table) at 1:100 dilution for neutrophils and LysoTracker Deep Red (key resources table) at 1:1000 dilution for lysosomes for 30 min. at 37°C in the dark. Cells were washed with MACS buffer and immediately imaged.

Fluorescence images were taken using either a Zeiss LSM710 or a Zeiss LSM980 inverted confocal laser scanning microscope equipped with a 63x(NA1.4) oil immersion objective. Detector wavelength cutoffs were set to minimize signal crosstalk between fluorophores.

### RNA isolation from pulmonary neutrophils

One hour after OMV i.a. injection, lungs were digested as described above. One animal had to be excluded, since the surgical time point has been missed. Neutrophils were isolated from single cell suspension using Streptavidin MicroBeads (Miltenyi Biotec, cat# 130-048-101) according to the manufacturer’s protocol. In brief, cells were counted and incubated with a biotinylated anti-mouse Ly-6G antibody (key resources table) for 20min. at 4°C. After washing, Streptavidin MicroBeads were added and incubated for 20min. at 4°C. After washing, a MidiMACS separator (Miltenyi Biotec, cat# 130-042-302) together with a LS column (Miltenyi Biotec, cat# 130-042-401) was used to isolate neutrophils and purity of positively selected neutrophils as well as unlabeled cells was assessed by flow cytometry (key resources table: Ly-6C PE-Cy7, CD11b APC). Neutrophils were pelleted, and supernatant was removed by pipetting. Neutrophils were snap frozen and stored at -80°C. RNA was isolated directly from frozen neutrophils pellets using Promega ReliaPrep RNA Cell Miniprep System (key resources table) according to the manufacturer’s protocol and quality was assessed using Bioanalyzer and RNA 6000 Nano Kit (RQN 5.8-9.6, median 7.55). Samples were snap frozen and stored at -80°C until further processing by the Next Generation Sequencing Platform, University of Bern, www.ngs.unibe.ch.

### RNA sequencing

RNA sequencing libraries were prepared using Lexogen CORALL total RNA-seq library kit according to the manufacturer’s protocol, which includes a rRNA depletion step. Sequencing was performed on an Illumina NovaSeq6000 SP flow cell, 2×50 cycles.

Alignment & quantification: The resulting fastq files were quality controlled using fastqc v0.11.9 (*Babraham Bioinformatics - FastQC A Quality Control Tool for High Throughput Sequence Data*, n.d.). Forward R1 reads were debarcoded by moving the first 12 nucleotides on the 5’ end to the name of the read via fastp v0.19.5 (Chen et al., 2018). The alignment was performed with STAR v2.7.10a_alpha_220818 (Dobin et al., 2013). First, a Genome index was generated from the mouse reference genome GRCm39 ENSEMBL version 108. Second, the reads were then aligned to the reference using STAR with default parameters. Gene read counts were quantified using featureCounts from subread v2.0.1 using the mm108 GTF annotation with default parameters (Liao et al., 2014).

Data visualization: The data from the read count matrix was normalized to reads per million (RPM) and log-transformed, x ➔ log(1+x). The resulting data was used for Principal Component Analysis (PCA), which was performed with the R function *prcomp* and visualized with a custom script using ggplot2 (Wickham, 2016).

Differential Gene expression: Differentially expressed genes were computed using the R package DESeq2 (Love et al., 2014).

Pathway enrichment analysis: The differentially expressed genes obtained from DESeq2 with an adjusted p-value below 0.01 were uploaded to Metascape for pathway analysis on 5^th^ December 2023 (Zhou et al., 2019).

Heatmaps: The lists of genes of the pathways of interest were obtained from genome.jp for the KEGG pathways and using the R function *gconvert* from the R package gprofiler2 otherwise (Kolberg & Raudvere, 2023). The log transformed data was used and the heatmaps were done with the heatmap.2 function of the R package gplots with the hierarchical cluster method “complete” using Pearson correlation distance (Warnes et al., 2022).

Volcano plot: The volcano plots were performed with the R package ggplot2 and a custom R script (Wickham, 2016).

### Statistical analysis

Descriptive statistics and statistical tests were performed using Prism software (key resources table) or R and Rstudio (key resources table). For differences between two groups a t-test was applied when data was normally distributed (parametric). Otherwise, a Wilcoxon rank sum test was used (non-parametric). For differences between more than 2 groups, a one-way ANOVA followed by a Tukey post-hoc test (parametric), or a Kruskal-Wallis test followed by a pairwise Wilcoxon rank-sum test with Benjamini-Hochberg (BH) correction (non-parametric) was used. For survival analyses, a log-rank test was used. No one-tailed tests were used. No method was used to predetermine experimental sample sizes. A linear model and the correlation between ATP and OD_600_ in Figure 2 and Figure 3 were computed using the R functions *lm* and *cor.test* with the following parameters:

stats::cor.test(*data*$*variable*ATP, *data*$*variable*growth, alternative = “two.sided”, method = “pearson”) summary(stats::lm(*variable*ATP ∼ *variable*growth, data = *data*))

P < 0.05 (p == padj, when correction for multiple testing was necessary) was considered significant in all statistical analyses unless stated otherwise in the figure legend. Significant differences are in *italic*.

## Acknowledgements

We are grateful for the technical and financial support of the Inselspital, the DBMR and our laboratory. A special thank goes to Dana Leuenberger and Isabel Büchi for their support with the animal experiments and microscopy. We also want to thank the Microscopy Imaging Center (MIC) of the University of Bern for their assistance in sample preparation and imaging on the electron microscope. We want to thank Prof. Dr. Dr. Fabio Grassi and his group for providing us with the pBAD28 and pHND10 plasmids. We also want to thank Prof. Dr. Siegfried Hapfelmeier, Dr. Olivier Schären and Prof. Dr. Torsten Seuberlich for their extremely helpful scientific support. We want to thank Prof. Dr. Andrew Macpherson supporting us with his expertise and providing us with the Keio collection. Finally, we want to thank Prof. Dr. Dr. Joel Zindel, Dr. Simone Zwicky and Dr. Felix Baier for their critical contributions and ideas to complete this project. Figures were created using BioRender.com.

## Data and code availability

RNA sequencing data will be made publicly available on the NCBI Sequence Read Archive (SRA) upon version of record (VOR) publication. In addition, we will then make publicly available all source data, metadata, and code to recreate the figures at BORIS portal of the University of Bern under a CC BY licence.

## Author contributions

Conceptualization: D.S. and G.B.; Methodology: D.S., G.B. and D.St.; Software: D.S., D.S.-T. and K.K.; Formal analysis: D.S., A.S, D.S.-T. and N.E.; Investigation D.S., A.S., S.M., N.E. and L.S.; Resources: G.B. and D.St.; Data curation: D.S., D.S.-T. and K.K.; Writing – original draft preparation: D.S., G.B., L.S., D. St., D.S.-T., K.K.; Visualisation: D.S., D.S.-T.; Supervision; G.B.; Project administration: D.S. and G.B.; Funding acquisition: G.B. and D.St.

---

**Figure 1-figure supplement 1.**
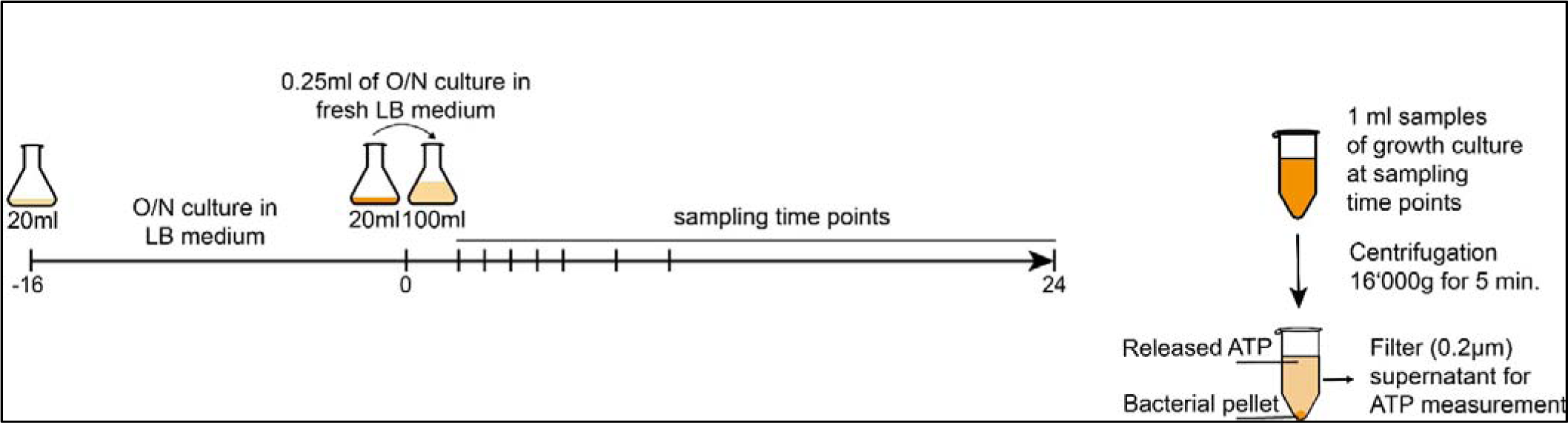
Experimental approach to measure released bacterial ATP and growth over time.

**Figure 2-figure supplement 1.**
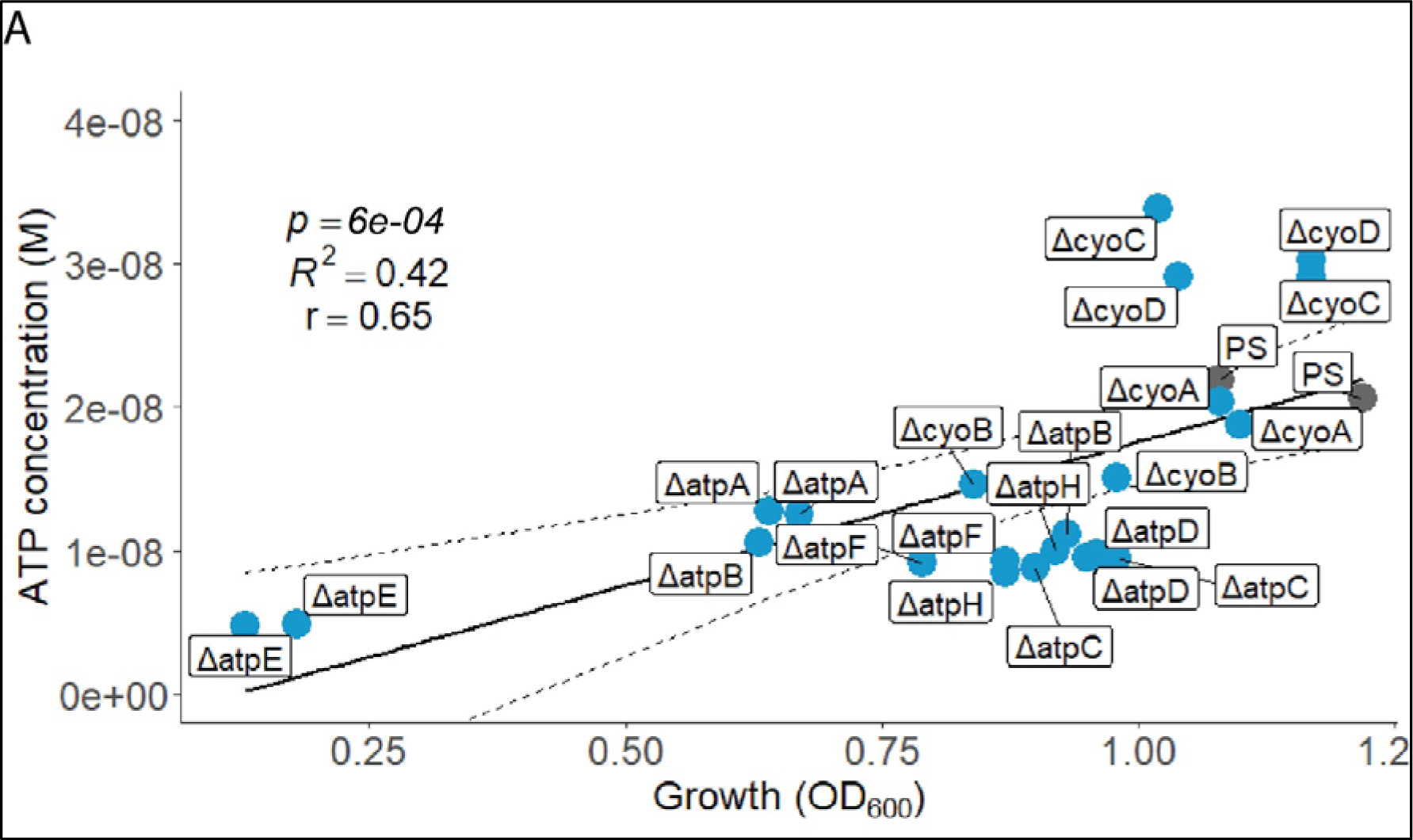
ATP concentration (M) and growth (OD_600_) at the individual peak of ATP release of all assessed cyo and atp mutants and the PS were plotted against each other. Pearson’s correlation (r) and coefficient of determination (*R*^2^) of the applied linear model are depicted. 95% confidence level is shown by the black dashed lines.

**Figure 4-figure supplement 1.**
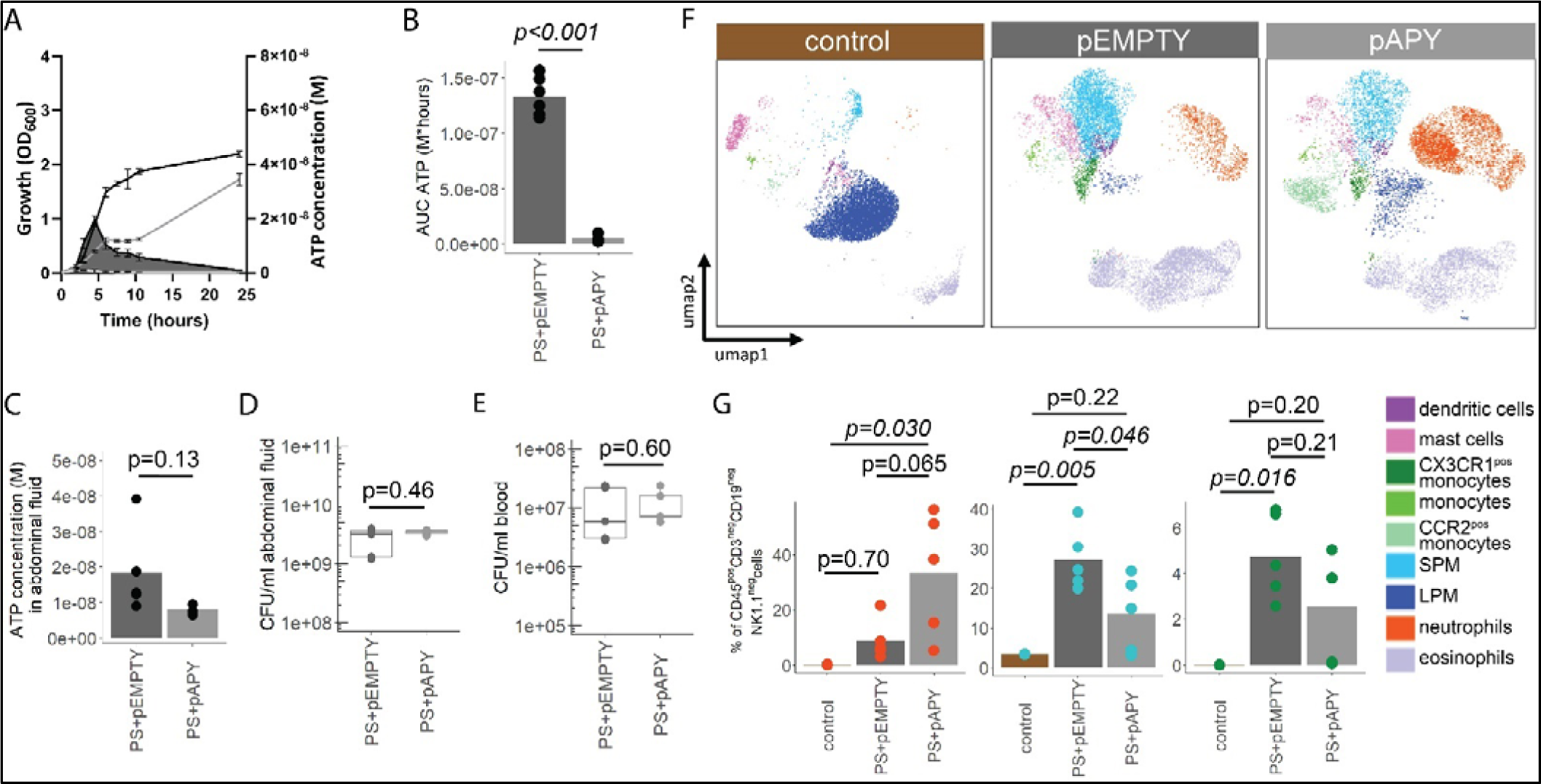
Immune cell characterization eight hours after i.a. injection of bacteria. **(A)** Measurement of released ATP (M) and growth (OD_600_) over time (hours) from PS+pEMPTY and PS+pAPY. n = 2 measurements of N = 3 independent bacteria cultures. Means and standard deviations are shown. **(B)** Area under the curve (AUC) of released ATP over time (M*hours) of the previously assessed bacteria (cumulative ATP). T-test, n = 2 measurements of N = 3 independent bacteria cultures. Means and individual values are shown. **(C)** Measurement of ATP (M) in abdominal fluid from mice eight hours after i.a. injection of bacteria. T-test, n = 5 animals per group of N = 2 independent experiments. Means and individual values are shown. **(D)** Quantitative assessment of colony forming units in abdominal fluid and **(E)** blood from mice eight hours after i.a. injection of bacteria. Wilcoxon rank sum test, n = 5 animals per group of N = 2 independent experiments. Means and individual values are shown. No growth for controls was detected. **(F)** Concatenated (n = 5 animals for each treatment group, n = 3 animals for control group of N = 2 independent experiments) and down-sampled images of immune cell populations characterized in the abdominal cavity eight hours after sham treatment or i.a. injection of bacteria. **G)** Abundance of neutrophils, small peritoneal macrophages (SPM) and CX3CR1^pos^ monocytes in abdominal fluid from mice eight hours after sham treatment or i.a. injection of bacteria. One-way ANOVA followed by Tukey post-hoc test, n = 5 animals for each treatment group, n = 3 animals for control group of N = 2 independent experiments. Means and individual values are shown.

**Figure 5-figure supplement 1.**
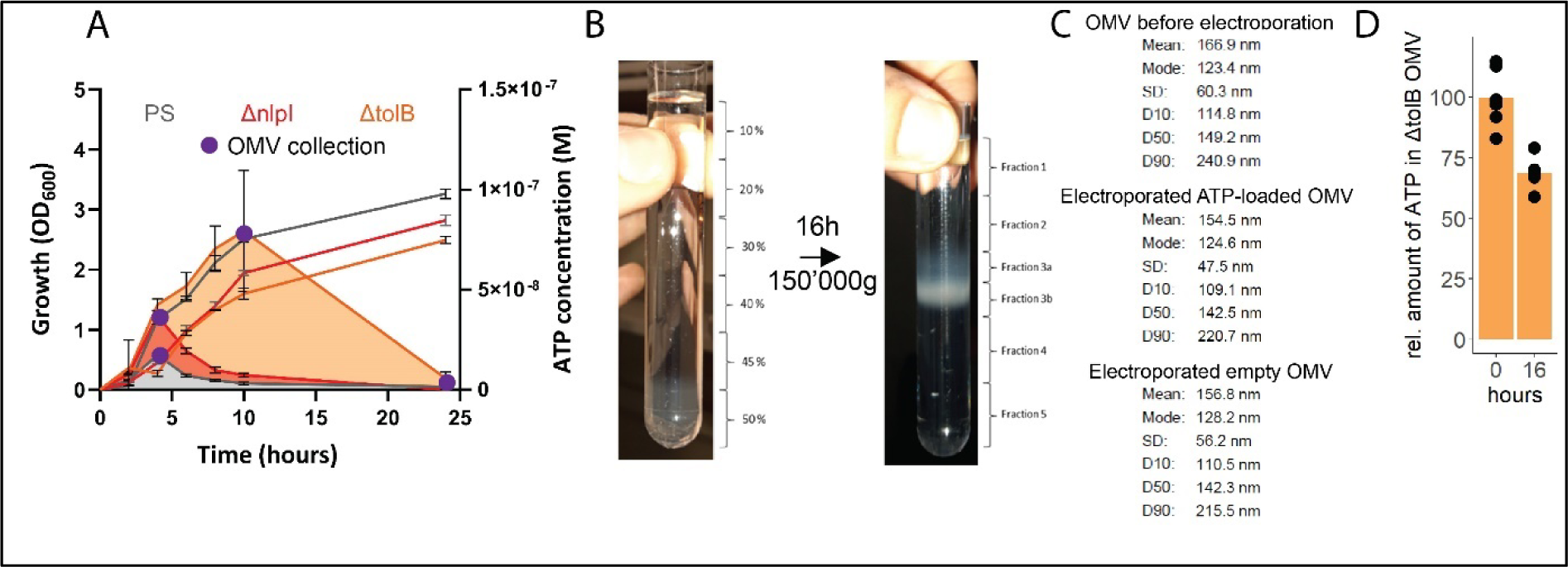
ATP measurement of the PS, Δ*nlpI* as well as Δ*tolB* and OMV collection and characterization. **(A)** Measurement of released ATP (M) and growth (OD_600_) over time (hours) from PS, Δ*nlpI* and Δ*tolB*. OMV collection time points are marked in purple. n = 2 measurements of N = 3 independent bacteria cultures. Means and standard deviations are shown. **(B)** OMV before and after density gradient ultracentrifugation for 16h at 150’000g. **(C)** Statistical parameters of OMV before electroporation as well as ATP-loaded and empty OMV after electroporation. **(D)** Relative quantification of ATP in OMV 16 hours at 4°C after electroporation (0h = 100%). n = 2 measurements of N = 3 independent experiments. Means and individual values are shown.

**Figure 6-figure supplement 1.**
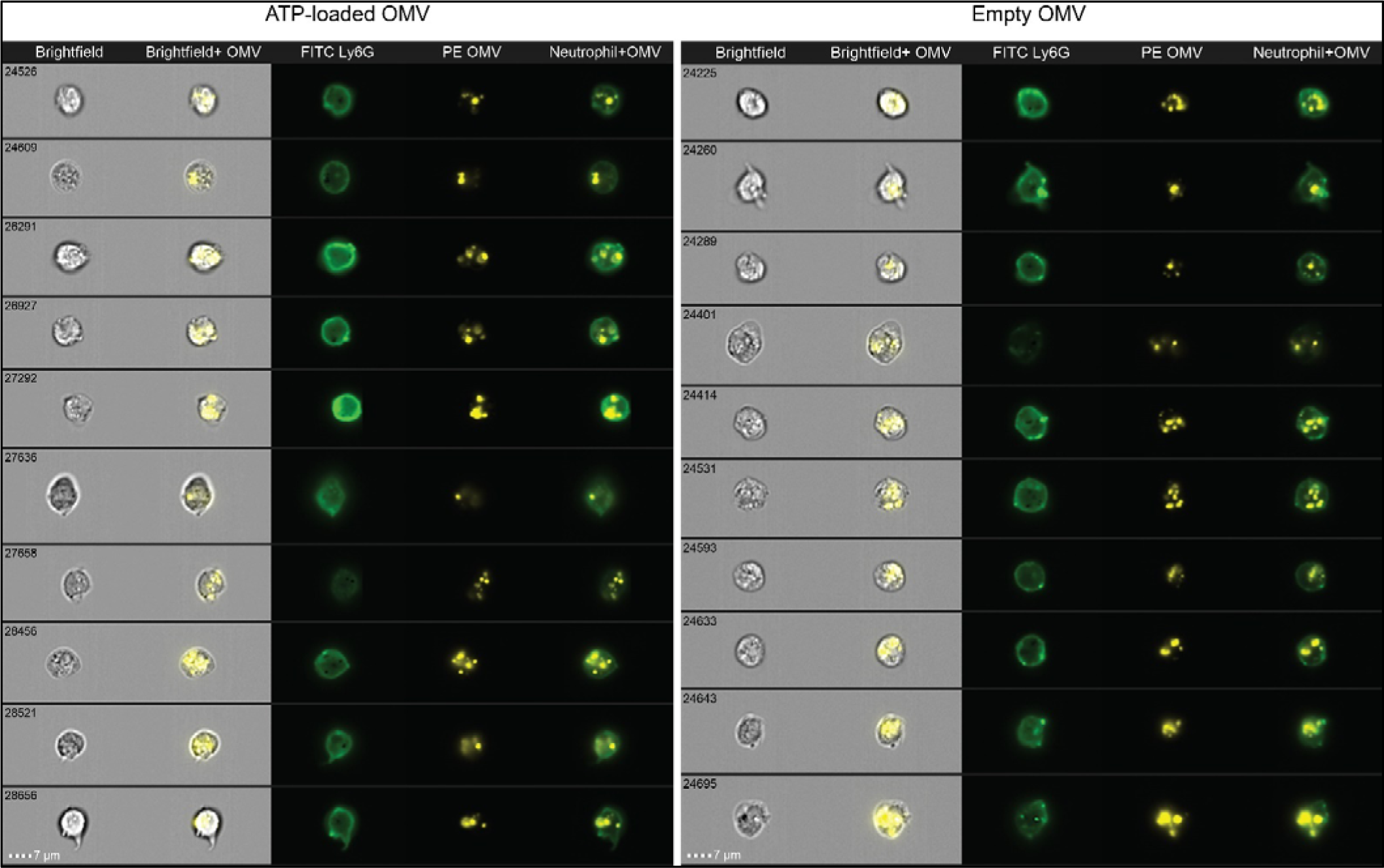
Uptake of OMV by neutrophils. Representative images of OMV uptake by neutrophils in the abdominal cavity one hour after i.a. injection additionally assessed using flow cytometry (Image Stream).

**Figure 6-figure supplement 2.**
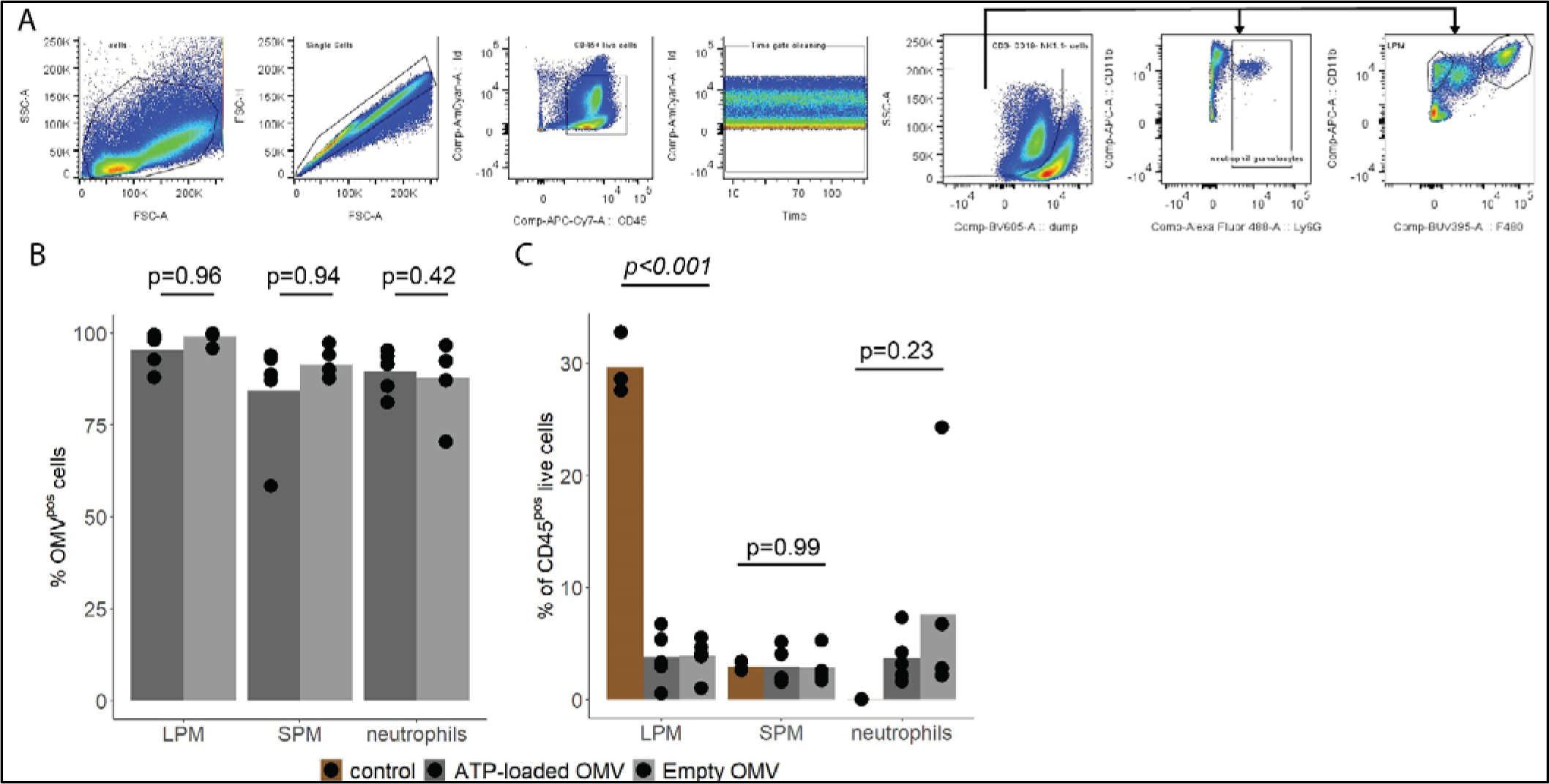
Characterization of local immune response in the abdominal cavity. **(A)** Gating strategy to identify large peritoneal macrophages (LPM), small peritoneal macrophages (SPM) and neutrophils in abdominal fluid. **(B)** Abundance of OMV^pos^ / (OMV^pos^+OMV^neg^) LPM, SPM and neutrophils one hour after i.a. injection of ATP-loaded or empty OMV. T-test with Benjamini-Hochberg correction, n = 5 animals per group of N = 2 independent experiments. Means and individual values are shown. **(C)** Abundance of LPM, SPM and neutrophils one hour after sham treatment or i.a. injection of either ATP-loaded or empty OMV. One-way ANOVA, n = 5 animals for each treatment group, n = 3 animals for control group of N = 2 independent experiments. Means and individual values are shown.

**Figure 6-figure supplement 3.**
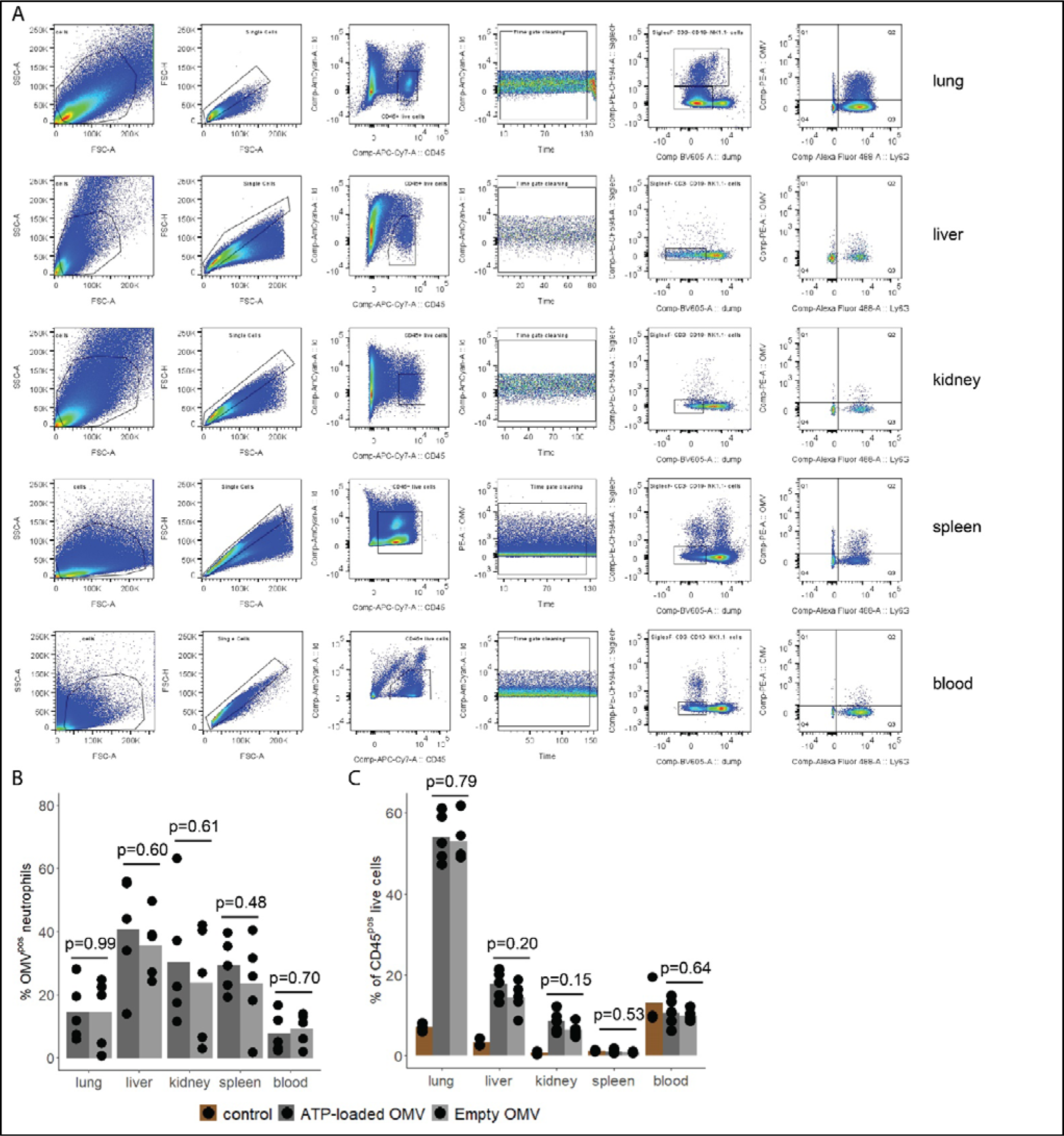
Assessment of OMV uptake by immune cells in remote organs. **(A)** Gating strategy to identify total OMV^pos^ cells and specifically OMV^pos^ neutrophils in blood and remote organs (lung, liver, kidney, and spleen) **(B)** Abundance of OMV^pos^ / (OMV^pos^+OMV^neg^) neutrophils one hour after i.a. injection of ATP-loaded or empty OMV. T-test, n = 5 animals per group of N = 2 independent experiments. Means and individual values are shown. **(C)** Neutrophils as percentage of all CD45^pos^ live cells one hour after sham treatment or i.a. injection of either ATP-loaded or empty OMV. T-test, n = 5 animals for each treatment group, n = 3 animals for control group of N = 2 independent experiments. Means and individual values are shown.

**Figure 6-figure supplement 4.**
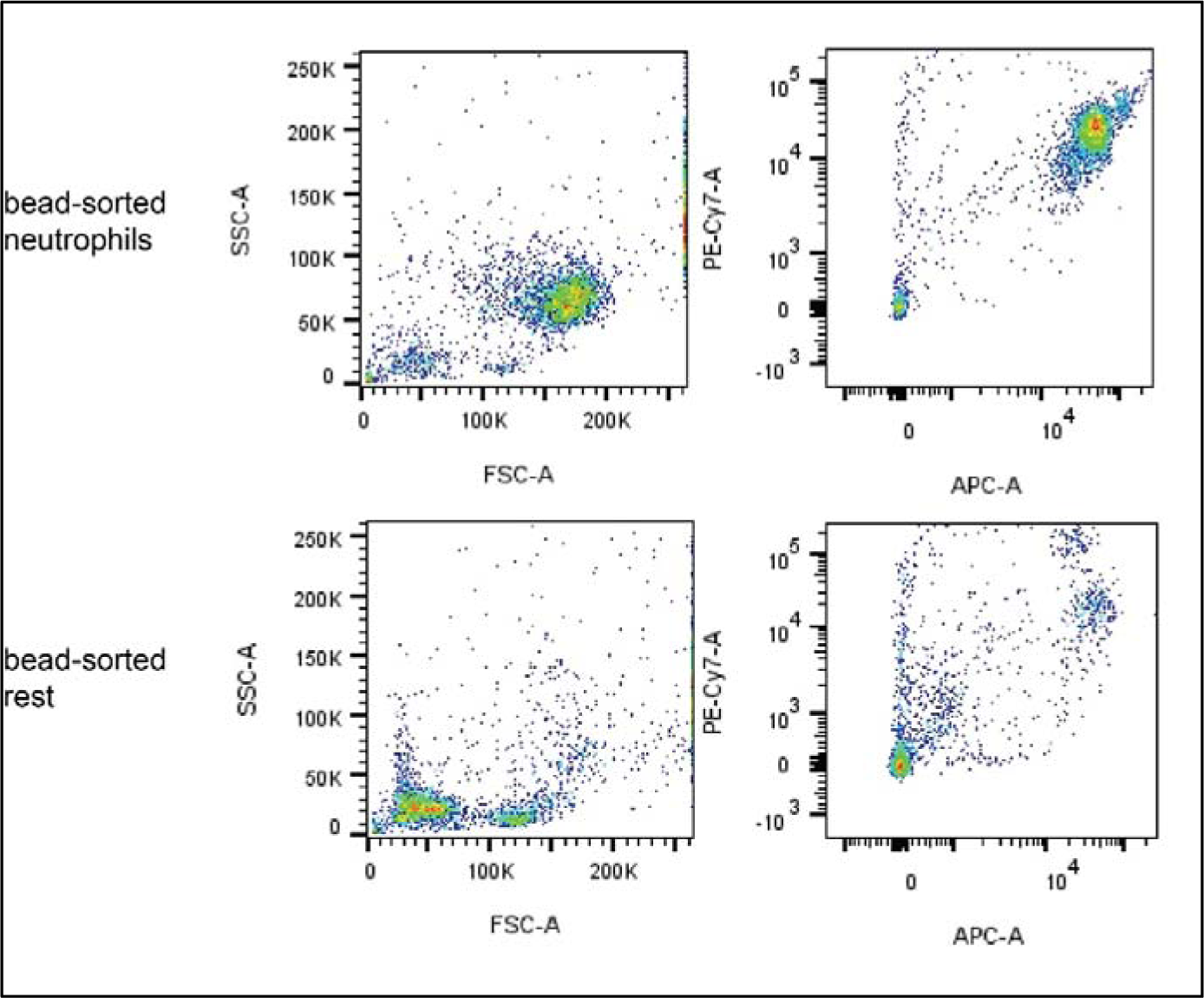
Assessment of the purity of bead-sorted pulmonary neutrophils. Pulmonary neutrophils were isolated one hour after i.a. injection of ATPγs-loaded or empty OMV, bead-sorted and assessed for purity by flow cytometry. A representative image is shown.

**Figure 6-figure supplement 5.**
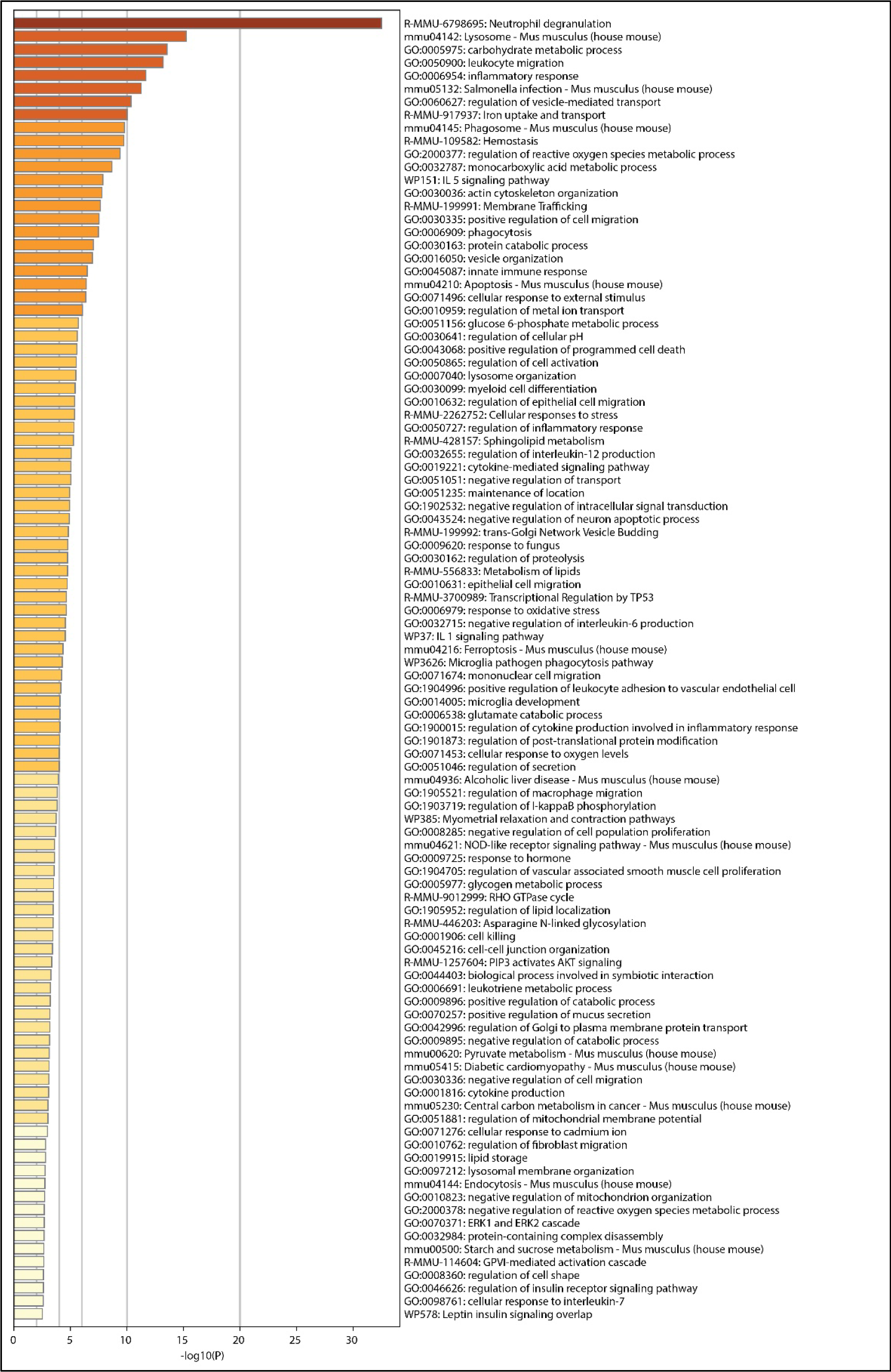
List of significantly different pathways after enrichment analysis of RNA sequencing results. Pulmonary neutrophils were isolated one hour after i.a. injection of ATPγs-loaded or empty OMV, bead-sorted and RNA sequencing was done. This resulted in these significantly different pathways between the groups after enrichment analysis. DESeq, n = 6 animals in the NE group, n = 5 animals in the NA group.

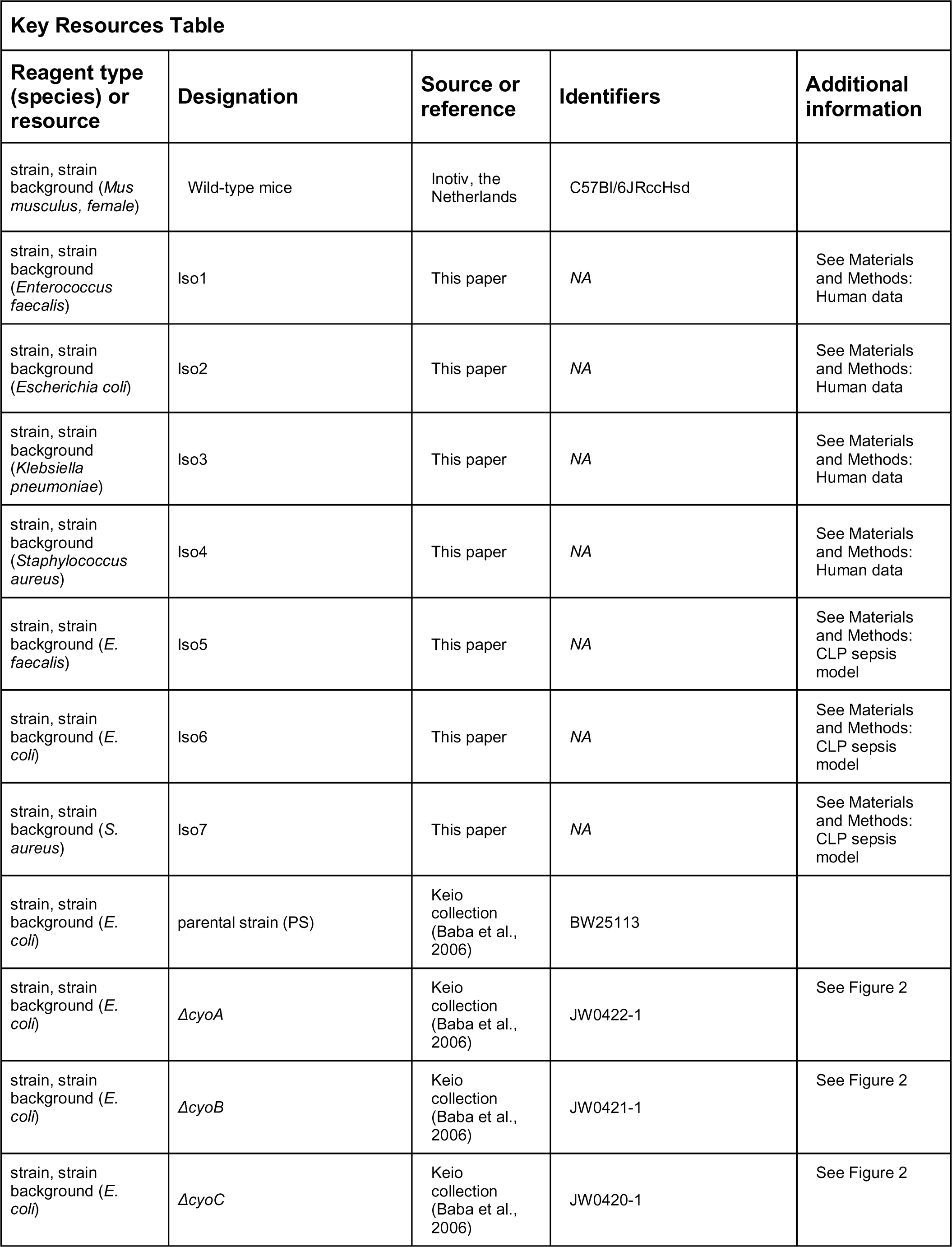
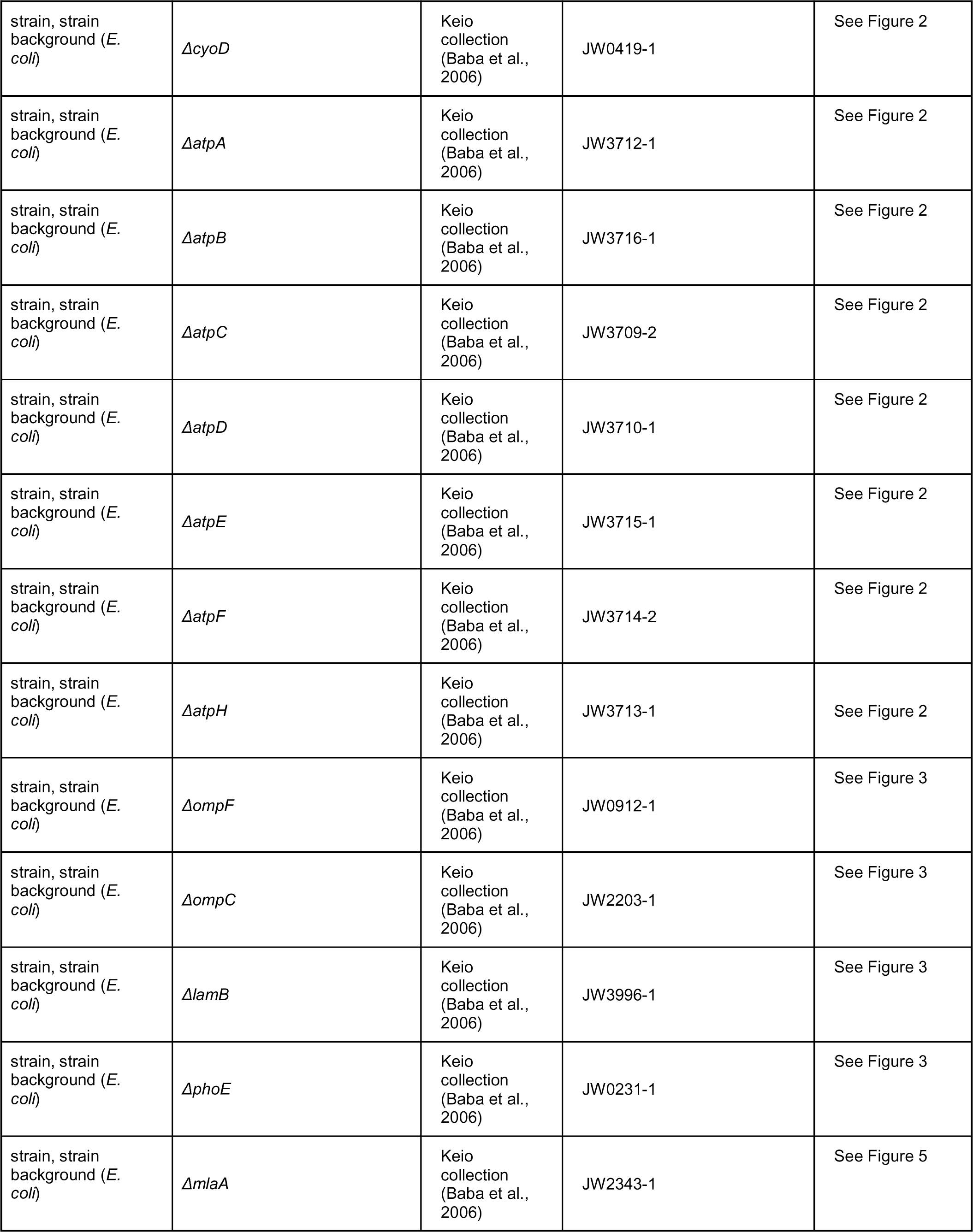
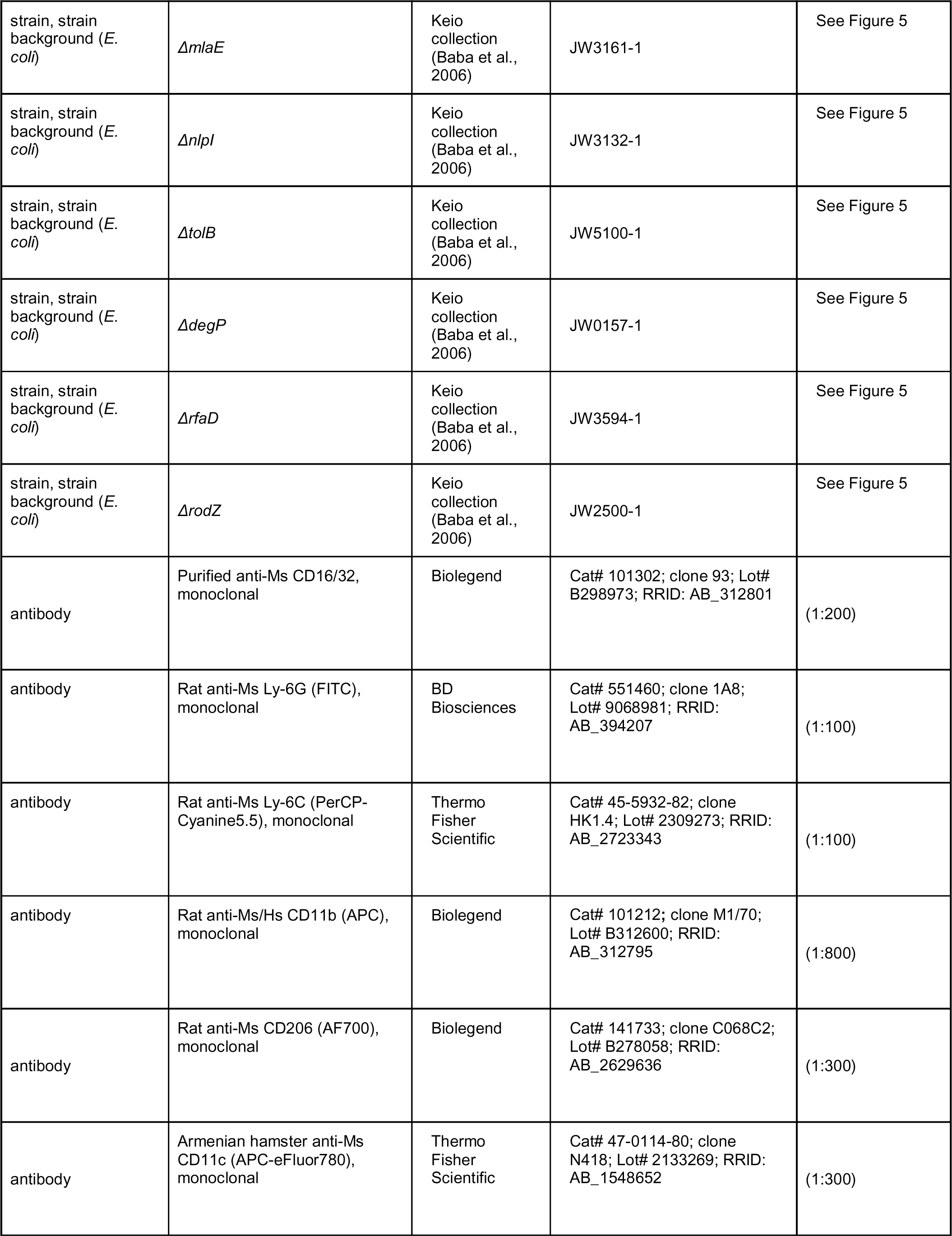
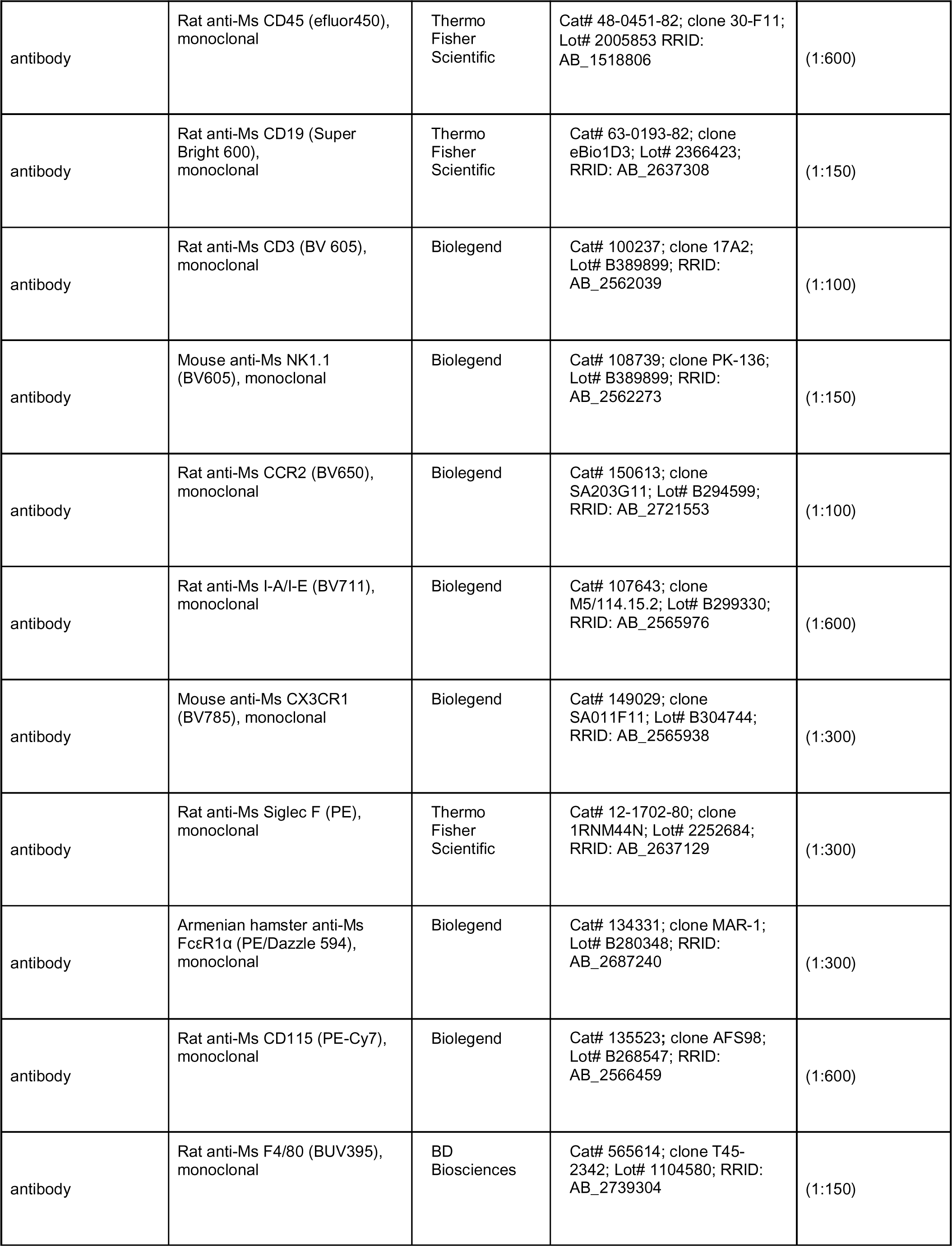
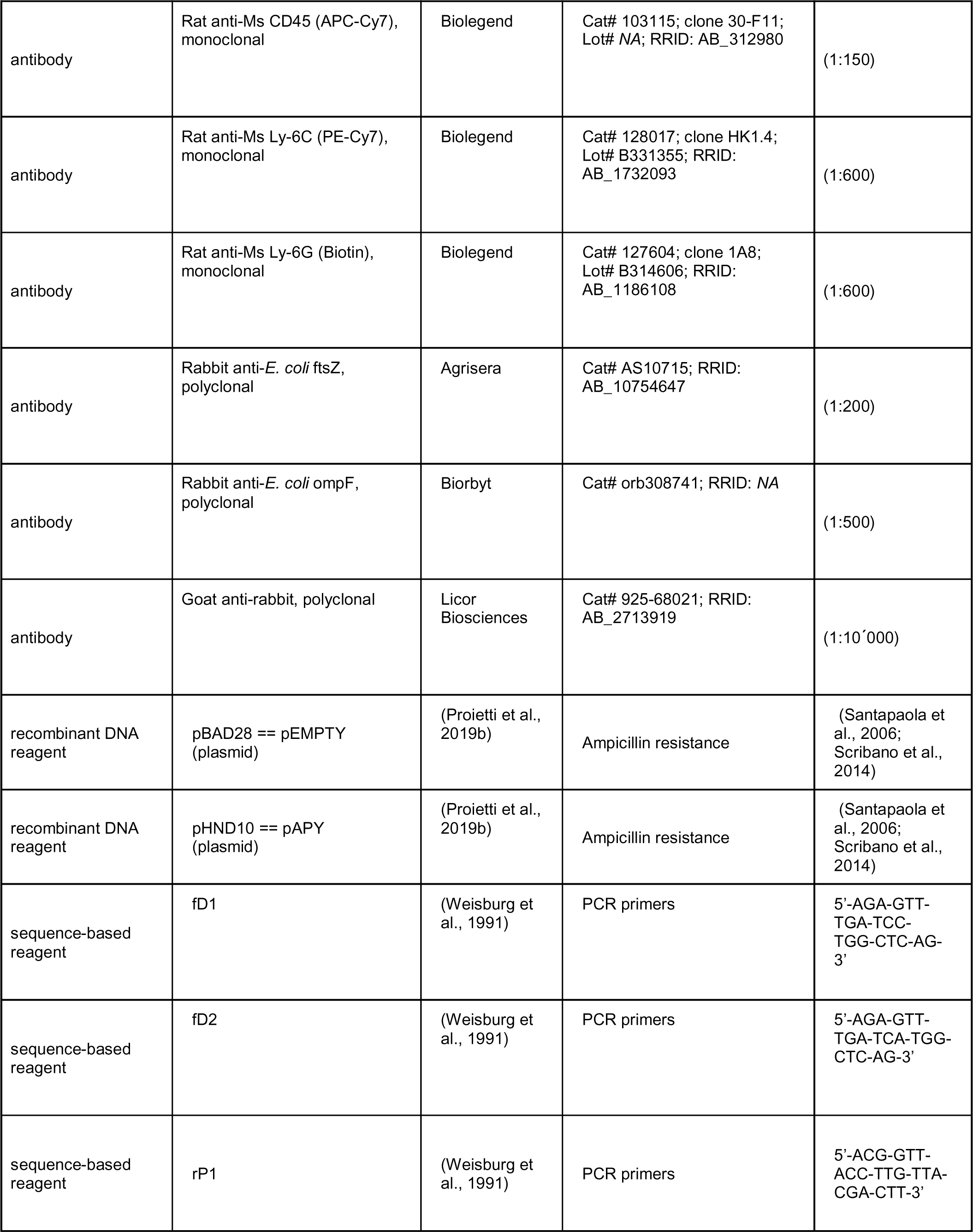
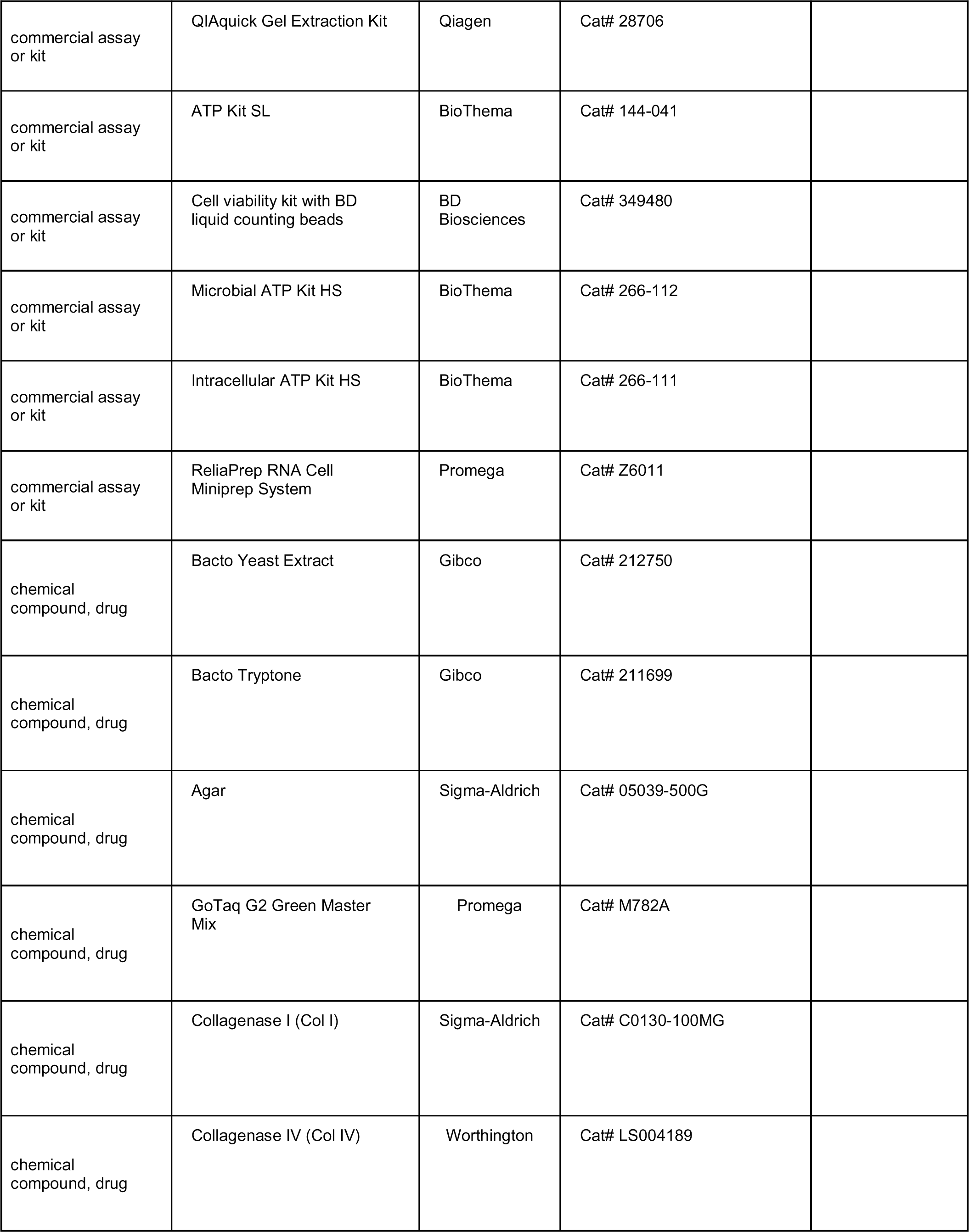
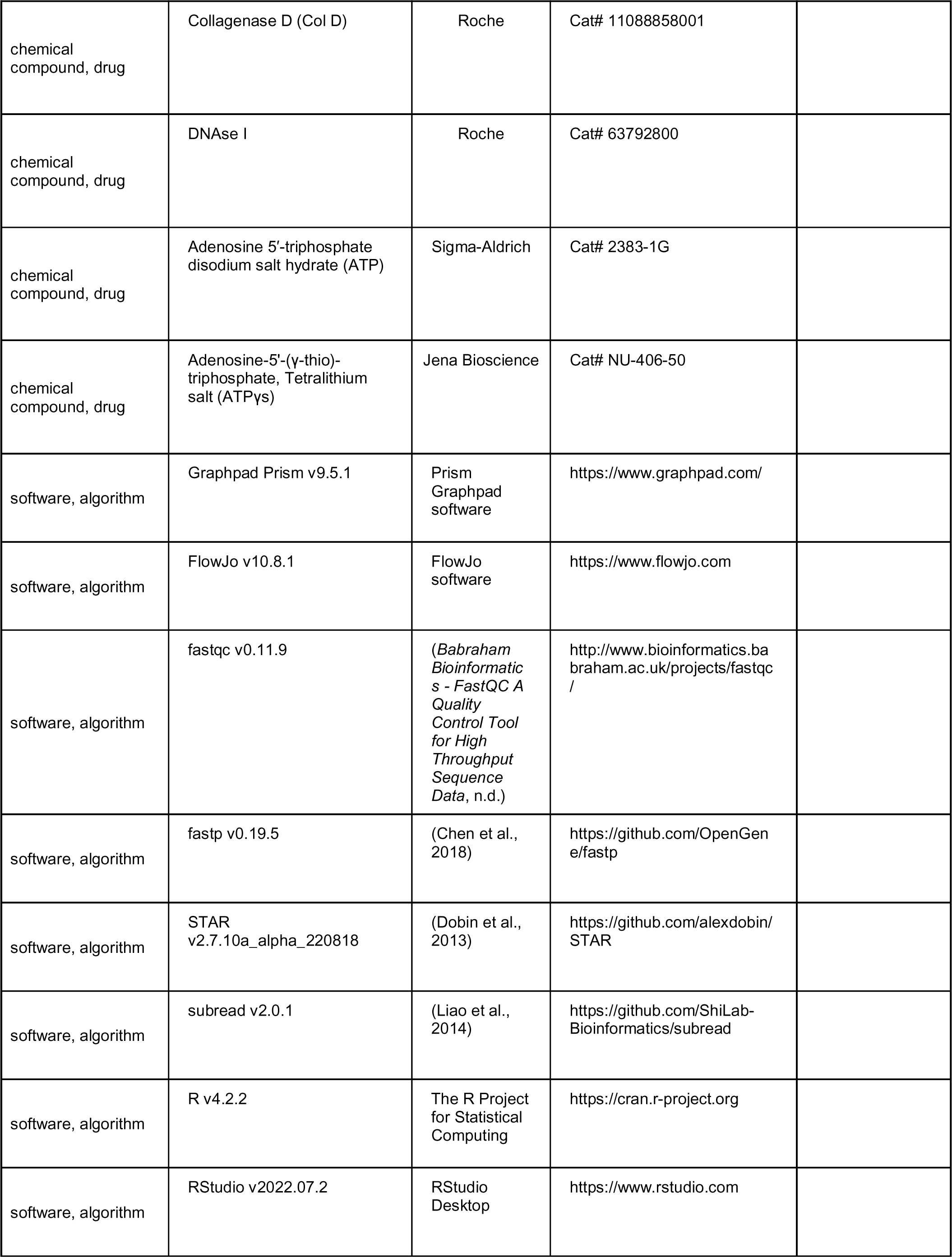
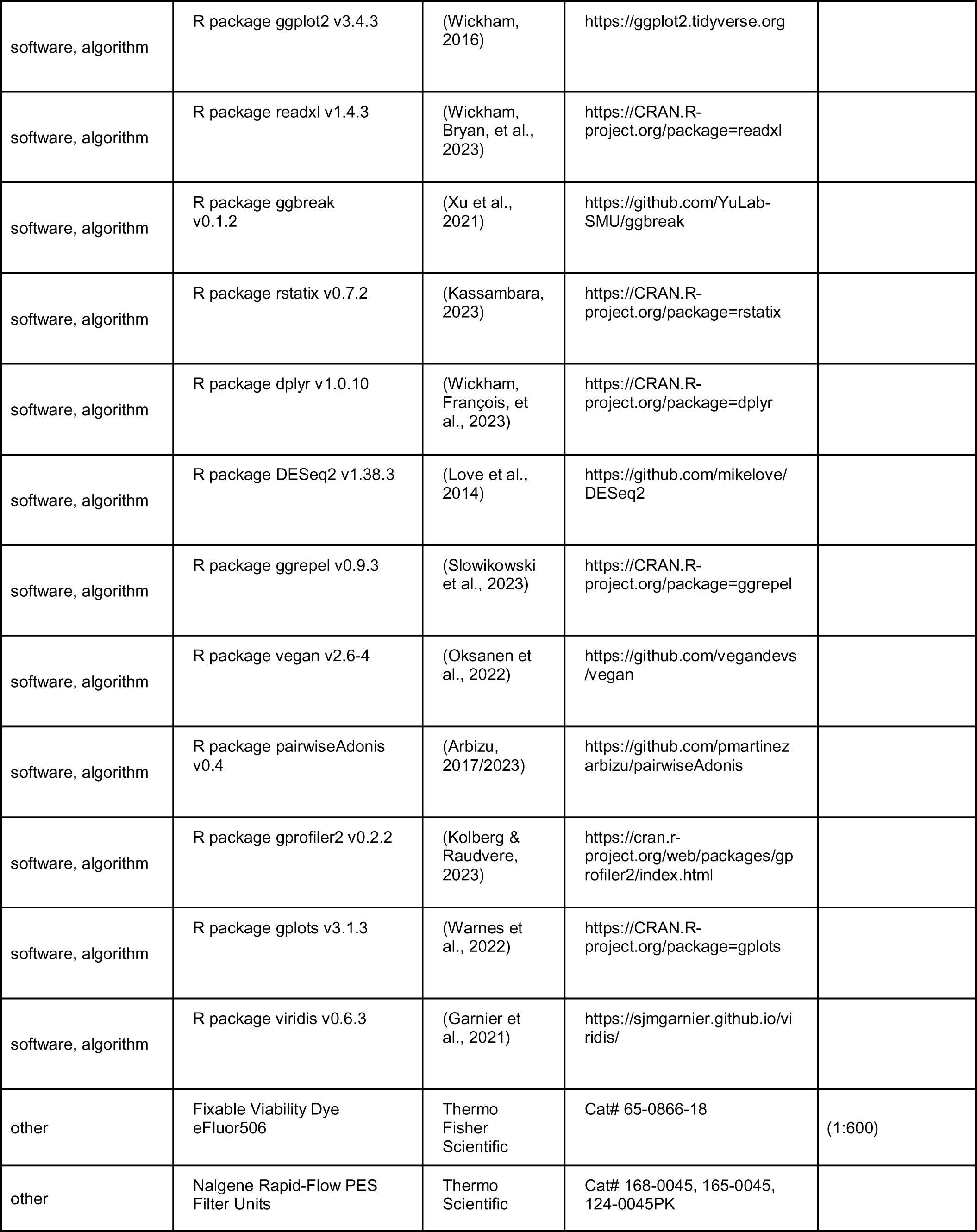
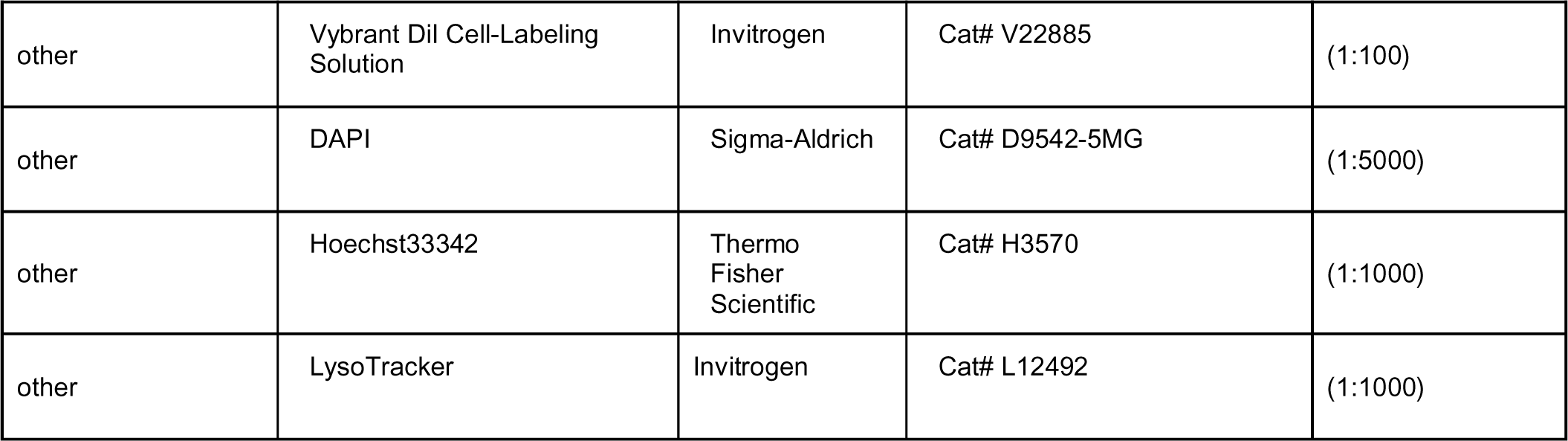

## References

Alvarez, C. L., Corradi, G., Lauri, N., Marginedas-Freixa, I., Leal Denis, M. F., Enrique, N., Mate, S. M., Milesi, V., Ostuni, M. A., Herlax, V., & Schwarzbaum, P. J. (2017). Dynamic regulation of extracellular ATP in Escherichia coli. Biochemical Journal, 474(8), 1395–1416. 10.1042/BCJ20160879

Araujo, M. E. G., Liebscher, G., Hess, M. W., & Huber, L. A. (2020). Lysosomal size matters. Traffic, 21(1), 60–75. 10.1111/tra.12714

Arbizu, P. M. (2023). pairwiseAdonis [R]. https://github.com/pmartinezarbizu/pairwiseAdonis (Original work published 2017)

Atarashi, K., Nishimura, J., Shima, T., Umesaki, Y., Yamamoto, M., Onoue, M., Yagita, H., Ishii, N., Evans, R., Honda, K., & Takeda, K. (2008). ATP drives lamina propria TH17 cell differentiation. Nature, 455(7214), 808–812. 10.1038/nature07240

Baba, T., Ara, T., Hasegawa, M., Takai, Y., Okumura, Y., Baba, M., Datsenko, K. A., Tomita, M., Wanner, B. L., & Mori, H. (2006). Construction of Escherichia coli K-12 in-frame, single-gene knockout mutants: The Keio collection. Molecular Systems Biology, 2(1), 2006.0008. 10.1038/msb4100050

Babraham Bioinformatics—FastQC A Quality Control tool for High Throughput Sequence Data. (n.d.). Retrieved January 17, 2024, from http://www.bioinformatics.babraham.ac.uk/projects/fastqc/

Baeza, N., & Mercade, E. (2021). Relationship Between Membrane Vesicles, Extracellular ATP and Biofilm Formation in Antarctic Gram-Negative Bacteria. Microbial Ecology, 81(3), 645–656. 10.1007/s00248-020-01614-6

Bielaszewska, M., Rüter, C., Bauwens, A., Greune, L., Jarosch, K.-A., Steil, D., Zhang, W., He, X., Lloubes, R., Fruth, A., Kim, K. S., Schmidt, M. A., Dobrindt, U., Mellmann, A., & Karch, H. (2017). Host cell interactions of outer membrane vesicle-associated virulence factors of enterohemorrhagic Escherichia coli O157: Intracellular delivery, trafficking and mechanisms of cell injury. PLOS Pathogens, 13(2), e1006159. 10.1371/journal.ppat.1006159

Bittel, M., Reichert, P., Sarfati, I., Dressel, A., Leikam, S., Uderhardt, S., Stolzer, I., Phu, T. A., Ng, M., Vu, N. K., Tenzer, S., Distler, U., Wirtz, S., Rothhammer, V., Neurath, M. F., Raffai, R. L., Günther, C., & Momma, S. (2021). Visualizing transfer of microbial biomolecules by outer membrane vesicles in microbe-host-communication in vivo. Journal of Extracellular Vesicles, 10(12). 10.1002/jev2.12159

Bitto, N. J., Chapman, R., Pidot, S., Costin, A., Lo, C., Choi, J., D’Cruze, T., Reynolds, E. C., Dashper, S. G., Turnbull, L., Whitchurch, C. B., Stinear, T. P., Stacey, K. J., & Ferrero, R. L. (2017). Bacterial membrane vesicles transport their DNA cargo into host cells. Scientific Reports, 7(1), 7072. 10.1038/s41598-017-07288-4

Borisov, V. B., Gennis, R. B., Hemp, J., & Verkhovsky, M. I. (2011). The cytochrome bd respiratory oxygen reductases. Biochimica et Biophysica Acta (BBA) - Bioenergetics, 1807(11), 1398–1413. 10.1016/j.bbabio.2011.06.016

Burnstock, G. (2020). Introduction to Purinergic Signaling. In P. Pelegrín (Ed.), Purinergic Signaling (Vol. 2041, pp. 1–15). Springer New York. 10.1007/978-1-4939-9717-6_1

Cao, M., Wang, G., & Xie, J. (2023). Immune dysregulation in sepsis: Experiences, lessons and perspectives. Cell Death Discovery, 9(1), 465. 10.1038/s41420-023-01766-7

Chen, S., Zhou, Y., Chen, Y., & Gu, J. (2018). fastp: An ultra-fast all-in-one FASTQ preprocessor. Bioinformatics (Oxford, England), 34(17), i884–i890. 10.1093/bioinformatics/bty560

Choi, U., & Lee, C.-R. (2019). Distinct Roles of Outer Membrane Porins in Antibiotic Resistance and Membrane Integrity in Escherichia coli. Frontiers in Microbiology, 10, 953. 10.3389/fmicb.2019.00953

Di Virgilio, F., Sarti, A. C., & Coutinho-Silva, R. (2020). Purinergic signaling, DAMPs, and inflammation. American Journal of Physiology-Cell Physiology, 318(5), C832–C835. 10.1152/ajpcell.00053.2020

Diekema, D. J., Hsueh, P.-R., Mendes, R. E., Pfaller, M. A., Rolston, K. V., Sader, H. S., & Jones, R. N. (2019). The Microbiology of Bloodstream Infection: 20-Year Trends from the SENTRY Antimicrobial Surveillance Program. Antimicrobial Agents and Chemotherapy, 63(7), e00355–19. 10.1128/AAC.00355-19

Ding, Q., Quah, S. Y., & Tan, K. S. (2016). Secreted adenosine triphosphate from Aggregatibacter actinomycetemcomitans triggers chemokine response. Molecular Oral Microbiology, 31(5), 423–434. 10.1111/omi.12143

Ding, Q., & Tan, K. S. (2016). The Danger Signal Extracellular ATP Is an Inducer of Fusobacterium nucleatum Biofilm Dispersal. Frontiers in Cellular and Infection Microbiology, 6. 10.3389/fcimb.2016.00155

Dobin, A., Davis, C. A., Schlesinger, F., Drenkow, J., Zaleski, C., Jha, S., Batut, P., Chaisson, M., & Gingeras, T. R. (2013). STAR: Ultrafast universal RNA-seq aligner. Bioinformatics (Oxford, England), 29(1), 15–21. 10.1093/bioinformatics/bts635

Dosch, M., Zindel, J., Jebbawi, F., Melin, N., Sanchez-Taltavull, D., Stroka, D., Candinas, D., & Beldi, G. (2019). Connexin-43-dependent ATP release mediates macrophage activation during sepsis. eLife, 8, e42670. 10.7554/eLife.42670

Eichelberger, K. R., & Goldman, W. E. (2020). Manipulating neutrophil degranulation as a bacterial virulence strategy. PLOS Pathogens, 16(12), e1009054. 10.1371/journal.ppat.1009054

Eltzschig, H. K., Sitkovsky, M. V., & Robson, S. C. (2012). Purinergic Signaling during Inflammation. New England Journal of Medicine, 367(24), 2322–2333. 10.1056/NEJMra1205750

Fu, S., Wang, Y., Xia, X., & Zheng, J. C. (2020). Exosome engineering: Current progress in cargo loading and targeted delivery. NanoImpact, 20, 100261. 10.1016/j.impact.2020.100261

Galber, C., Carissimi, S., Baracca, A., & Giorgio, V. (2021). The ATP Synthase Deficiency in Human Diseases. Life, 11(4), Article 4. 10.3390/life11040325

Garnier, S., Ross, N., Rudis, boB, Filipovic-Pierucci, A., Galili, T., timelyportfolio, Greenwell, B., Sievert, C., Harris, D. J., & Chen, J. J. (2021). sjmgarnier/viridis: Viridis 0.6.0 (pre-CRAN release) (v0.6.0pre) [Computer software]. Zenodo. 10.5281/zenodo.4679424

Gheinani, A. H., Vögeli, M., Baumgartner, U., Vassella, E., Draeger, A., Burkhard, F. C., & Monastyrskaya, K. (2018). Improved isolation strategies to increase the yield and purity of human urinary exosomes for biomarker discovery. Scientific Reports, 8(1), 3945. 10.1038/s41598-018-22142-x

Ghosn, E. E. B., Cassado, A. A., Govoni, G. R., Fukuhara, T., Yang, Y., Monack, D. M., Bortoluci, K. R., Almeida, S. R., Herzenberg, L. A., & Herzenberg, L. A. (2010). Two physically, functionally, and developmentally distinct peritoneal macrophage subsets. Proceedings of the National Academy of Sciences, 107(6), 2568–2573. 10.1073/pnas.0915000107

Grauel, A., Kägi, J., Rasmussen, T., Makarchuk, I., Oppermann, S., Moumbock, A. F. A., Wohlwend, D., Müller, R., Melin, F., Günther, S., Hellwig, P., Böttcher, B., & Friedrich, T. (2021). Structure of Escherichia coli cytochrome bd-II type oxidase with bound aurachin D. Nature Communications, 12(1), 6498. 10.1038/s41467-021-26835-2

Hironaka, I., Iwase, T., Sugimoto, S., Okuda, K., Tajima, A., Yanaga, K., & Mizunoe, Y. (2013). Glucose Triggers ATP Secretion from Bacteria in a Growth-Phase-Dependent Manner. Applied and Environmental Microbiology, 79(7), 2328–2335. 10.1128/AEM.03871-12

Ihssen, J., Jovanovic, N., Sirec, T., & Spitz, U. (2021). Real-time monitoring of extracellular ATP in bacterial cultures using thermostable luciferase. PLOS ONE, 16(1), e0244200. 10.1371/journal.pone.0244200

Ivančić, V., Mastali, M., Percy, N., Gornbein, J., Babbitt, J. T., Li, Y., Landaw, E. M., Bruckner, D. A., Churchill, B. M., & Haake, D. A. (2008). Rapid Antimicrobial Susceptibility Determination of Uropathogens in Clinical Urine Specimens by Use of ATP Bioluminescence. Journal of Clinical Microbiology, 46(4), 1213–1219. 10.1128/JCM.02036-07

Iwase, T., Shinji, H., Tajima, A., Sato, F., Tamura, T., Iwamoto, T., Yoneda, M., & Mizunoe, Y. (2010). Isolation and Identification of ATP-Secreting Bacteria from Mice and Humans. Journal of Clinical Microbiology, 48(5), 1949–1951. 10.1128/JCM.01941-09

Jang, S. C., Kim, S. R., Yoon, Y. J., Park, K.-S., Kim, J. H., Lee, J., Kim, O. Y., Choi, E.-J., Kim, D.-K., Choi, D.-S., Kim, Y.-K., Park, J., Di Vizio, D., & Gho, Y. S. (2015). In vivo Kinetic Biodistribution of Nano-Sized Outer Membrane Vesicles Derived from Bacteria. Small, 11(4), 456–461. 10.1002/smll.201401803

Junger, W. G. (2011). Immune cell regulation by autocrine purinergic signalling. Nature Reviews Immunology, 11(3), 201–212. 10.1038/nri2938

Juodeikis, R., & Carding, S. R. (2022). Outer Membrane Vesicles: Biogenesis, Functions, and Issues. Microbiology and Molecular Biology Reviews, 86(4), e00032–22. 10.1128/mmbr.00032-22

Kassambara, A. (2023). rstatix: Pipe-Friendly Framework for Basic Statistical Tests (0.7.2) [Computer software]. https://cran.r-project.org/web/packages/rstatix/index.html

Kim, J. H., Yoon, Y. J., Lee, J., Choi, E.-J., Yi, N., Park, K.-S., Park, J., Lötvall, J., Kim, Y.-K., & Gho, Y. S. (2013). Outer Membrane Vesicles Derived from Escherichia coli Up-Regulate Expression of Endothelial Cell Adhesion Molecules In Vitro and In Vivo. PLoS ONE, 8(3), e59276. 10.1371/journal.pone.0059276

Koch, A. L. (1959). DEATH OF BACTERIA IN GROWING CULTURE. Journal of Bacteriology, 77(5), 623–629. 10.1128/jb.77.5.623-629.1959

Kolberg, L., & Raudvere, U. (2023). gprofiler2: Interface to the “g:Profiler” Toolset (0.2.2) [Computer software]. https://cran.r-project.org/web/packages/gprofiler2/index.html

Kulp, A., & Kuehn, M. J. (2010). Biological Functions and Biogenesis of Secreted Bacterial Outer Membrane Vesicles. Annual Review of Microbiology, 64(1), 163–184. 10.1146/annurev.micro.091208.073413

Ledderose, C., Bao, Y., Kondo, Y., Fakhari, M., Slubowski, C., Zhang, J., & Junger, W. G. (2016). Purinergic Signaling and the Immune Response in Sepsis: A Review. Clinical Therapeutics, 38(5), 1054–1065. 10.1016/j.clinthera.2016.04.002

Leduc, M., & Van Heijenoort, J. (1980). Autolysis of Escherichia coli. Journal of Bacteriology, 142(1), 52–59. 10.1128/jb.142.1.52-59.1980

Lee, E.-Y., Bang, J. Y., Park, G. W., Choi, D.-S., Kang, J. S., Kim, H.-J., Park, K.-S., Lee, J.-O., Kim, Y.-K., Kwon, K.-H., Kim, K.-P., & Gho, Y. S. (2007). Global proteomic profiling of native outer membrane vesicles derived from Escherichia coli . PROTEOMICS, 7(17), 3143–3153. 10.1002/pmic.200700196

Lee, J., Yoon, Y. J., Kim, J. H., Dinh, N. T. H., Go, G., Tae, S., Park, K.-S., Park, H. T., Lee, C., Roh, T.-Y., Di Vizio, D., & Gho, Y. S. (2018). Outer Membrane Vesicles Derived From Escherichia coli Regulate Neutrophil Migration by Induction of Endothelial IL-8. Frontiers in Microbiology, 9, 2268. 10.3389/fmicb.2018.02268

Leive, L. (1968). Studies on the Permeability Change Produced in Coliform Bacteria by Ethylenediaminetetraacetate. Journal of Biological Chemistry, 243(9), 2373–2380. 10.1016/S0021-9258(18)93484-8

Lennaárd, A. J., Mamand, D. R., Wiklander, R. J., EL Andaloussi, S., & Wiklander, O. P. B. (2021). Optimised Electroporation for Loading of Extracellular Vesicles with Doxorubicin. Pharmaceutics, 14(1), 38. 10.3390/pharmaceutics14010038

Liao, Y., Smyth, G. K., & Shi, W. (2014). featureCounts: An efficient general purpose program for assigning sequence reads to genomic features. Bioinformatics (Oxford, England), 30(7), 923–930. 10.1093/bioinformatics/btt656

Love, M. I., Huber, W., & Anders, S. (2014). Moderated estimation of fold change and dispersion for RNA-seq data with DESeq2. Genome Biology, 15(12), 550. 10.1186/s13059-014-0550-8

Lundin, A. (2000). Use of firefly luciferase in atp-related assays of biomass, enzymes, and metabolites. In Methods in Enzymology (Vol. 305, pp. 346–370). Elsevier. 10.1016/S0076-6879(00)05499-9

McBroom, A. J., Johnson, A. P., Vemulapalli, S., & Kuehn, M. J. (2006). Outer Membrane Vesicle Production by Escherichia coli Is Independent of Membrane Instability. Journal of Bacteriology, 188(15), 5385–5392. 10.1128/JB.00498-06

Mempin, R., Tran, H., Chen, C., Gong, H., Kim Ho, K., & Lu, S. (2013). Release of extracellular ATP by bacteria during growth. BMC Microbiology, 13(1), 301. 10.1186/1471-2180-13-301

Michel, L. V., & Gaborski, T. (2022). Outer membrane vesicles as molecular biomarkers for Gram-negative sepsis: Taking advantage of nature’s perfect packages. Journal of Biological Chemistry, 298(10), 102483. 10.1016/j.jbc.2022.102483

Mureșan, M. G., Balmoș, I. A., Badea, I., & Santini, A. (2018). Abdominal Sepsis: An Update. The Journal of Critical Care Medicine, 4(4), 120–125. 10.2478/jccm-2018-0023

Oksanen, J., Simpson, G. L., Blanchet, F. G., Kindt, R., Legendre, P., Minchin, P. R., O’Hara, R. B., Solymos, P., Stevens, M. H. H., Szoecs, E., Wagner, H., Barbour, M., Bedward, M., Bolker, B., Borcard, D., Carvalho, G., Chirico, M., Caceres, M. D., Durand, S., … Weedon, J. (2022). vegan: Community Ecology Package (2.6-4) [Computer software]. https://cran.r-project.org/web/packages/vegan/index.html

Orench-Rivera, N., & Kuehn, M. J. (2016). Environmentally controlled bacterial vesicle-mediated export: Environmentally controlled bacterial vesicle-mediated export. Cellular Microbiology, 18(11), 1525–1536. 10.1111/cmi.12676

Park, K.-S., Choi, K.-H., Kim, Y.-S., Hong, B. S., Kim, O. Y., Kim, J. H., Yoon, C. M., Koh, G.-Y., Kim, Y.-K., & Gho, Y. S. (2010). Outer Membrane Vesicles Derived from Escherichia coli Induce Systemic Inflammatory Response Syndrome. PLoS ONE, 5(6), e11334. 10.1371/journal.pone.0011334

Park, K.-S., Lee, J., Jang, S. C., Kim, S. R., Jang, M. H., Lötvall, J., Kim, Y.-K., & Gho, Y. S. (2013). Pulmonary Inflammation Induced by Bacteria-Free Outer Membrane Vesicles from Pseudomonas aeruginosa. American Journal of Respiratory Cell and Molecular Biology, 49(4), 637–645. 10.1165/rcmb.2012-0370OC

Pérez-Cruz, C., Carrión, O., Delgado, L., Martinez, G., López-Iglesias, C., & Mercade, E. (2013). New Type of Outer Membrane Vesicle Produced by the Gram-Negative Bacterium Shewanella vesiculosa M7T: Implications for DNA Content. Applied and Environmental Microbiology, 79(6), 1874–1881. 10.1128/AEM.03657-12

Pérez-Cruz, C., Delgado, L., López-Iglesias, C., & Mercade, E. (2015). Outer-Inner Membrane Vesicles Naturally Secreted by Gram-Negative Pathogenic Bacteria. PLOS ONE, 10(1), e0116896. 10.1371/journal.pone.0116896

Perruzza, L., Gargari, G., Proietti, M., Fosso, B., D’Erchia, A. M., Faliti, C. E., Rezzonico-Jost, T., Scribano, D., Mauri, L., Colombo, D., Pellegrini, G., Moregola, A., Mooser, C., Pesole, G., Nicoletti, M., Norata, G. D., Geuking, M. B., McCoy, K. D., Guglielmetti, S., & Grassi, F. (2017). T Follicular Helper Cells Promote a Beneficial Gut Ecosystem for Host Metabolic Homeostasis by Sensing Microbiota-Derived Extracellular ATP. Cell Reports, 18(11), 2566–2575. 10.1016/j.celrep.2017.02.061

Proietti, M., Perruzza, L., Scribano, D., Pellegrini, G., D’Antuono, R., Strati, F., Raffaelli, M., Gonzalez, S. F., Thelen, M., Hardt, W.-D., Slack, E., Nicoletti, M., & Grassi, F. (2019a). ATP released by intestinal bacteria limits the generation of protective IgA against enteropathogens. Nature Communications, 10(1), Article 1. 10.1038/s41467-018-08156-z

Proietti, M., Perruzza, L., Scribano, D., Pellegrini, G., D’Antuono, R., Strati, F., Raffaelli, M., Gonzalez, S. F., Thelen, M., Hardt, W.-D., Slack, E., Nicoletti, M., & Grassi, F. (2019b). ATP released by intestinal bacteria limits the generation of protective IgA against enteropathogens. Nature Communications, 10(1), Article 1. 10.1038/s41467-018-08156-z

Qureshi, O. S., Paramasivam, A., Yu, J. C. H., & Murrell-Lagnado, R. D. (2007). Regulation of P2X4 receptors by lysosomal targeting, glycan protection and exocytosis. Journal of Cell Science, 120(21), 3838–3849. 10.1242/jcs.010348

Ray, A., & Dittel, B. N. (2010). Isolation of Mouse Peritoneal Cavity Cells. Journal of Visualized ExperimentsflJ: JoVE, 35. 10.3791/1488

Reimer, S. L., Beniac, D. R., Hiebert, S. L., Booth, T. F., Chong, P. M., Westmacott, G. R., Zhanel, G. G., & Bay, D. C. (2021). Comparative Analysis of Outer Membrane Vesicle Isolation Methods With an Escherichia coli tolA Mutant Reveals a Hypervesiculating Phenotype With Outer-Inner Membrane Vesicle Content. Frontiers in Microbiology, 12, 628801. 10.3389/fmicb.2021.628801

Reinhart, K., Daniels, R., Kissoon, N., Machado, F. R., Schachter, R. D., & Finfer, S. (2017). Recognizing Sepsis as a Global Health Priority—A WHO Resolution. New England Journal of Medicine, 377(5), 414–417. 10.1056/NEJMp1707170

Rittirsch, D., Huber-Lang, M. S., Flierl, M. A., & Ward, P. A. (2009). Immunodesign of experimental sepsis by cecal ligation and puncture. Nature Protocols, 4(1), Article 1. 10.1038/nprot.2008.214

Robinson, L. E., & Murrell-Lagnado, R. D. (2013). The trafficking and targeting of P2X receptors. Frontiers in Cellular Neuroscience, 7. 10.3389/fncel.2013.00233

Rudd, K. E., Johnson, S. C., Agesa, K. M., Shackelford, K. A., Tsoi, D., Kievlan, D. R., Colombara, D. V., Ikuta, K. S., Kissoon, N., Finfer, S., Fleischmann-Struzek, C., Machado, F. R., Reinhart, K. K., Rowan, K., Seymour, C. W., Watson, R. S., West, T. E., Marinho, F., Hay, S. I., … Naghavi, M. (2020). Global, regional, and national sepsis incidence and mortality, 1990–2017: Analysis for the Global Burden of Disease Study. The Lancet, 395(10219), 200–211. 10.1016/S0140-6736(19)32989-7

Salm, L., Shim, R., Noskovicova, N., & Kubes, P. (2023). Gata6+ large peritoneal macrophages: An evolutionarily conserved sentinel and effector system for infection and injury. Trends in Immunology, 44(2), 129–145. 10.1016/j.it.2022.12.002

Santapaola, D., Del Chierico, F., Petrucca, A., Uzzau, S., Casalino, M., Colonna, B., Sessa, R., Berlutti, F., & Nicoletti, M. (2006). Apyrase, the product of the virulence plasmid-encoded phoN2 (apy) gene of Shigella flexneri, is necessary for proper unipolar IcsA localization and for efficient intercellular spread. Journal of Bacteriology, 188(4), 1620–1627. 10.1128/JB.188.4.1620-1627.2006

Sartorio, M. G., Pardue, E. J., Feldman, M. F., & Haurat, M. F. (2021). Bacterial Outer Membrane Vesicles: From Discovery to Applications. Annual Review of Microbiology, 75(1), 609–630. 10.1146/annurev-micro-052821-031444

Schwechheimer, C., & Kuehn, M. J. (2015). Outer-membrane vesicles from Gram-negative bacteria: Biogenesis and functions. Nature Reviews Microbiology, 13(10), 605–619. 10.1038/nrmicro3525

Scribano, D., Petrucca, A., Pompili, M., Ambrosi, C., Bruni, E., Zagaglia, C., Prosseda, G., Nencioni, L., Casalino, M., Polticelli, F., & Nicoletti, M. (2014). Polar Localization of PhoN2, a Periplasmic Virulence-Associated Factor of Shigella flexneri, Is Required for Proper IcsA Exposition at the Old Bacterial Pole. PLOS ONE, 9(2), e90230. 10.1371/journal.pone.0090230

Slowikowski, K., Schep, A., Hughes, S., Dang, T. K., Lukauskas, S., Irisson, J.-O., Kamvar, Z. N., Ryan, T., Christophe, D., Hiroaki, Y., Gramme, P., Abdol, A. M., Barrett, M., Cannoodt, R., Krassowski, M., Chirico, M., Aphalo, P., & Barton, F. (2023). ggrepel: Automatically Position Non-Overlapping Text Labels with “ggplot2” (0.9.4) [Computer software]. https://cran.r-project.org/web/packages/ggrepel/index.html

Spari, D., & Beldi, G. (2020). Extracellular ATP as an Inter-Kingdom Signaling Molecule: Release Mechanisms by Bacteria and Its Implication on the Host. International Journal of Molecular Sciences, 21(15), Article 15. 10.3390/ijms21155590

Takaki, K., Tahara, Y. O., Nakamichi, N., Hasegawa, Y., Shintani, M., Ohkuma, M., Miyata, M., Futamata, H., & Tashiro, Y. (2020). Multilamellar and Multivesicular Outer Membrane Vesicles Produced by a Buttiauxella agrestis tolB Mutant. Applied and Environmental Microbiology, 86(20), e01131–20. 10.1128/AEM.01131-20

Toyofuku, M., Nomura, N., & Eberl, L. (2019). Types and origins of bacterial membrane vesicles. Nature Reviews Microbiology, 17(1), 13–24. 10.1038/s41579-018-0112-2

Tsubaki, M., Hori, H., & Mogi, T. (2000). Probing molecular structure of dioxygen reduction site of bacterial quinol oxidases through ligand binding to the redox metal centers. Journal of Inorganic Biochemistry, 82(1–4), 19–25. 10.1016/S0162-0134(00)00140-9

Turnbull, L., Toyofuku, M., Hynen, A. L., Kurosawa, M., Pessi, G., Petty, N. K., Osvath, S. R., Cárcamo-Oyarce, G., Gloag, E. S., Shimoni, R., Omasits, U., Ito, S., Yap, X., Monahan, L. G., Cavaliere, R., Ahrens, C. H., Charles, I. G., Nomura, N., Eberl, L., & Whitchurch, C. B. (2016). Explosive cell lysis as a mechanism for the biogenesis of bacterial membrane vesicles and biofilms. Nature Communications, 7(1), 11220. 10.1038/ncomms11220

Turner, L., Praszkier, J., Hutton, M. L., Steer, D., Ramm, G., Kaparakis-Liaskos, M., & Ferrero, R. L. (2015). Increased Outer Membrane Vesicle Formation in a Helicobacter pylori tolB Mutant. Helicobacter, 20(4), 269–283. 10.1111/hel.12196

Van Gassen, S., Callebaut, B., Van Helden, M. J., Lambrecht, B. N., Demeester, P., Dhaene, T., & Saeys, Y. (2015). FlowSOM: Using self-organizing maps for visualization and interpretation of cytometry data. Cytometry Part A, 87(7), 636–645. 10.1002/cyto.a.22625

Vega-Pérez, A., Villarrubia, L. H., Godio, C., Gutiérrez-González, A., Feo-Lucas, L., Ferriz, M., Martínez-Puente, N., Alcaín, J., Mora, A., Sabio, G., López-Bravo, M., & Ardavín, C. (2021). Resident macrophage-dependent immune cell scaffolds drive anti-bacterial defense in the peritoneal cavity. Immunity, 54(11), 2578–2594.e5. 10.1016/j.immuni.2021.10.007

Wang, D.-H., Jia, H.-M., Zheng, X., Xi, X.-M., Zheng, Y., & Li, W.-X. (2024). Attributable mortality of ARDS among critically ill patients with sepsis: A multicenter, retrospective cohort study. BMC Pulmonary Medicine, 24(1), 110. 10.1186/s12890-024-02913-1

Warnes, G. R., Bolker, B., Bonebakker, L., Gentleman, R., Huber, W., Liaw, A., Lumley, T., Maechler, M., Magnusson, A., Moeller, S., Schwartz, M., Venables, B., & Galili, T. (2022). gplots: Various R Programming Tools for Plotting Data (3.1.3) [Computer software]. https://cran.r-project.org/web/packages/gplots/index.html

Weisburg, W. G., Barns, S. M., Pelletier, D. A., & Lane, D. J. (1991). 16S ribosomal DNA amplification for phylogenetic study. Journal of Bacteriology, 173(2), 697–703. 10.1128/jb.173.2.697-703.1991

Wickham, H. (2016). Ggplot2. Springer International Publishing. 10.1007/978-3-319-24277-4

Wickham, H., Bryan, J., Posit, attribution), P. (Copyright holder of all R. code and all C. code without explicit copyright, code), M. K. (Author of included R., code), K. V. (Author of included libxls, code), C. L. (Author of included libxls, code), B. C. (Author of included libxls, code), D. H. (Author of included libxls, & code), E. M. (Author of included libxls. (2023). readxl: Read Excel Files (1.4.3) [Computer software]. https://cran.r-project.org/web/packages/readxl/index.html

Wickham, H., François, R., Henry, L., Müller, K., Vaughan, D., Software, P., & PBC. (2023). dplyr: A Grammar of Data Manipulation (1.1.4) [Computer software]. https://cran.r-project.org/web/packages/dplyr/index.html

Xu, S., Chen, M., Feng, T., Zhan, L., Zhou, L., & Yu, G. (2021). Use ggbreak to Effectively Utilize Plotting Space to Deal With Large Datasets and Outliers. Frontiers in Genetics, 12. https://www.frontiersin.org/articles/10.3389/fgene.2021.774846

Yamamoto, N., Nakahigashi, K., Nakamichi, T., Yoshino, M., Takai, Y., Touda, Y., Furubayashi, A., Kinjyo, S., Dose, H., Hasegawa, M., Datsenko, K. A., Nakayashiki, T., Tomita, M., Wanner, B. L., & Mori, H. (2009). Update on the Keio collection of Escherichia coli single-gene deletion mutants. Molecular Systems Biology, 5(1), 335. 10.1038/msb.2009.92

Yegutkin, G. G., Mikhailov, A., Samburski, S. S., & Jalkanen, S. (2006). The Detection of Micromolar Pericellular ATP Pool on Lymphocyte Surface by Using Lymphoid Ecto-Adenylate Kinase as Intrinsic ATP SensorLlV. Molecular Biology of the Cell, 17.

Zhou, Y., Zhou, B., Pache, L., Chang, M., Khodabakhshi, A. H., Tanaseichuk, O., Benner, C., & Chanda, S. K. (2019). Metascape provides a biologist-oriented resource for the analysis of systems-level datasets. Nature Communications, 10(1), 1523. 10.1038/s41467-019-09234-6

Zindel, J., Peiseler, M., Hossain, M., Deppermann, C., Lee, W. Y., Haenni, B., Zuber, B., Deniset, J. F., Surewaard, B. G. J., Candinas, D., & Kubes, P. (2021). Primordial GATA6 macrophages function as extravascular platelets in sterile injury. Science, 371(6533), eabe0595. 10.1126/science.abe0595

